# Tracking tau fibrillogenesis and consequent primary phagocytosis of neurons mediated by microglia in a living tauopathy model

**DOI:** 10.1101/2020.11.04.368977

**Authors:** Hiroyuki Takuwa, Asumi Orihara, Yuhei Takado, Takuya Urushihata, Masafumi Shimojo, Ai Ishikawa, Manami Takahashi, Anna M. Barron, Maiko Ono, Jun Maeda, Kazuto Masamoto, Hiroyasu Akatsu, Aviva M. Tolkovsky, Bin Ji, Yutaka Tomita, Hiroshi Ito, Ming-Rong Zhang, Michel Goedert, Maria Grazia Spillantini, Virginia M. -Y. Lee, John Q. Trojanowski, Taketoshi Maehara, Tetsuya Suhara, Naruhiko Sahara, Makoto Higuchi

## Abstract

Fibrillary tau pathologies have been implicated in Alzheimer’s and allied neurodegenerative diseases, while mechanisms by which neurons bearing tau tangles die remain enigmatic. To address this issue, we pursued tau and related key pathologies macroscopically by PET and MRI and microscopically by intravital two-photon laser optics. Time-course macroscopic assays of tau transgenic mice demonstrated intimate associations of tau deposition and increase of an inflammatory microglial marker, translocator protein (TSPO), with regional brain atrophy. Longitudinal microscopy of these mice revealed a rapid turnover of tau lesions resulting from continuous generation of new tau aggregates followed by loss of neurons and their fibrillar contents. This technology also allowed the capturing of the disappearance of tangle-bearing neurons several days after being engulfed by activated microglia. Notably, a therapeutic TSPO ligand profoundly suppressed the mobility and phagocytotic activity of microglia and improved neuronal survival in this model, supporting the involvement of primary phagocytosis of viable neurons by microglia in tau-primed neuronal death. Finally, partial depletion of microglia revealed roles of immune factors, MFG-E8 and C1q, as ‘eat-me’ signals for an immediate attraction of phagocytic microglia towards the elimination of tangle-loaded neurons.

## INTRODUCTION

Microtubule-associated protein tau (MAPT) is a constituent of axonal cytoskeletons in normally functioning neurons, while self-assembly of microtubule-unbound tau into fibrils is a pathological hallmark in diverse neuropathological disorders (collectively referred to as tauopathies) exemplified by Alzheimer’s disease (AD) ^1^. The discovery of *MAPT* mutations causative of familial tauopathy termed frontotemporal dementia and parkinsonism linked to chromosome 17 *MAPT* (FTDP-17-*MAPT*) provided compelling evidence for mechanistic links between tau fibrillogenesis and neuronal deteriorations ^2^. However, molecular processes mediating tau-induced neurotoxicity are yet to be clarified. Oligomeric tau species may provoke neuronal death in a cell-autonomous fashion ^3,4^, which could be circumvented by packaging misfolded tau molecules in a tangle of dense fibrils. In the meantime, soluble tau escaping from this packaging machinery may disseminate in the extracellular matrix to evoke deleterious gliosis that can accordingly attack neurons with and without tangles ^5,6^.

Indeed, inflammatory microgliosis has been documented to precede neuronal loss and to accelerate neurodegenerative tau pathologies in a mouse model transgenic for a human tau isoform with the FTDP-17-*MAPT* mutations ^6^. In these strains, inhibitions of neuroinflammatory processes by immunosuppressants and anti-inflammatory agents resulted in attenuation of tau accumulations and neuronal death ^6^, and deficiencies and overexpressions of microglial receptors, including fractalkine receptor and triggering receptor expressed on myeloid cells 2, lead to profound modulations of tau aggregation and neurodegeneration ^7–9^. Moreover, treatment of these animals with lipopolysaccharide (LPS), a stimulant of toll-like receptor-4 and downstream inflammatory signaling mainly in microglia, was reported to exacerbate neurodegenerative damages ^7,10^. Despite this various evidence for the loss of neurons carrying tau aggregates elicited by aggressive microglia, little is known about whether microglia-mediated neurotoxicity rather than self-destruction of neurons is a dominant factor governing tau-provoked neuronal death. In addition, how detrimental microglial cells kill neurons in this pathological condition is still unclarified. Such specific modes of microglial actions affecting neuronal survival may include microglial production of reactive oxygen species giving lethal damage to neurons, while elimination of live neurons could be mediated by microglial phagocytosis ^11^.

We have recently established a bimodal *in-vivo* imaging system capable of fibrillary tau deposits in the mouse brain with cellular to regional/subregional resolutions, which takes advantage of a specific chemical probe applicable to two-photon laser fluorescence microscopy and positron emission tomography (PET) ^12**Error!Referencesourcenotfound.**^. The intravital microscopic technology is of particular utility when combined with visualization of fluorescence proteins distinctly expressed in neurons and microglia, as it allows us to track the chronological sequence of tau deposition, interaction with microglia, and death at an individual neuronal level.

## RESULT

### Intimate associations among tau accumulations, inflammatory microgliosis and brain atrophy and their dissociated suppressions by transgene inactivation in a mouse model

We employed a transgenic mouse strain dubbed rTg4510 ^13^ for a longitudinal pursuit of tau-triggered pathological events. This model expresses a P301L mutant human tau isoform specifically in neurons under control of the Tet-Off system, and develops neuronal tau inclusions resembling neurofibrillary tangles (NFTs) in tauopathies, in concurrence with glial activations and neuronal loss ^13**Error!Referencesourcenotfound.**^. Each animal underwent time-course PET imaging of tau and neuroinflammation and volumetric MRI, as described elsewhere ^14,15^. A bimodal optical and radiological imaging agent for tau aggregates, PBB3 ^12, 14, 15**Error! Reference source not found.**^, was radiolabeled with ^11^C, and was applied to the PET visualization of tau tangles. Likewise, a PET assessment of inflammatory microglial responses was carried out with AC-5216, which is a ^11^C-labeled probe for 18-kDa translocator protein (TSPO), a biomarker for deleterious microglia ^6,16,17^.

Longitudinal macroscopic assays captured the evolution of tau pathology in close association with progressive activation of TSPO-positive microglia and reduction of the neocortical and hippocampal volumes (Figure 1). It is noteworthy that the accumulation of PBB3-positive tau deposits appeared to decelerate beyond the age of 6 months, in distinction from neuroinflammation and regional atrophy linearly proportional to age (Figure 1). The non-linear advancement of tau deposition implies tangle formation and loss of tangle-burdened neurons in parallel, and the sum of these two events may promote the linear progress in microgliosis and volume loss.

**Figure 1.**
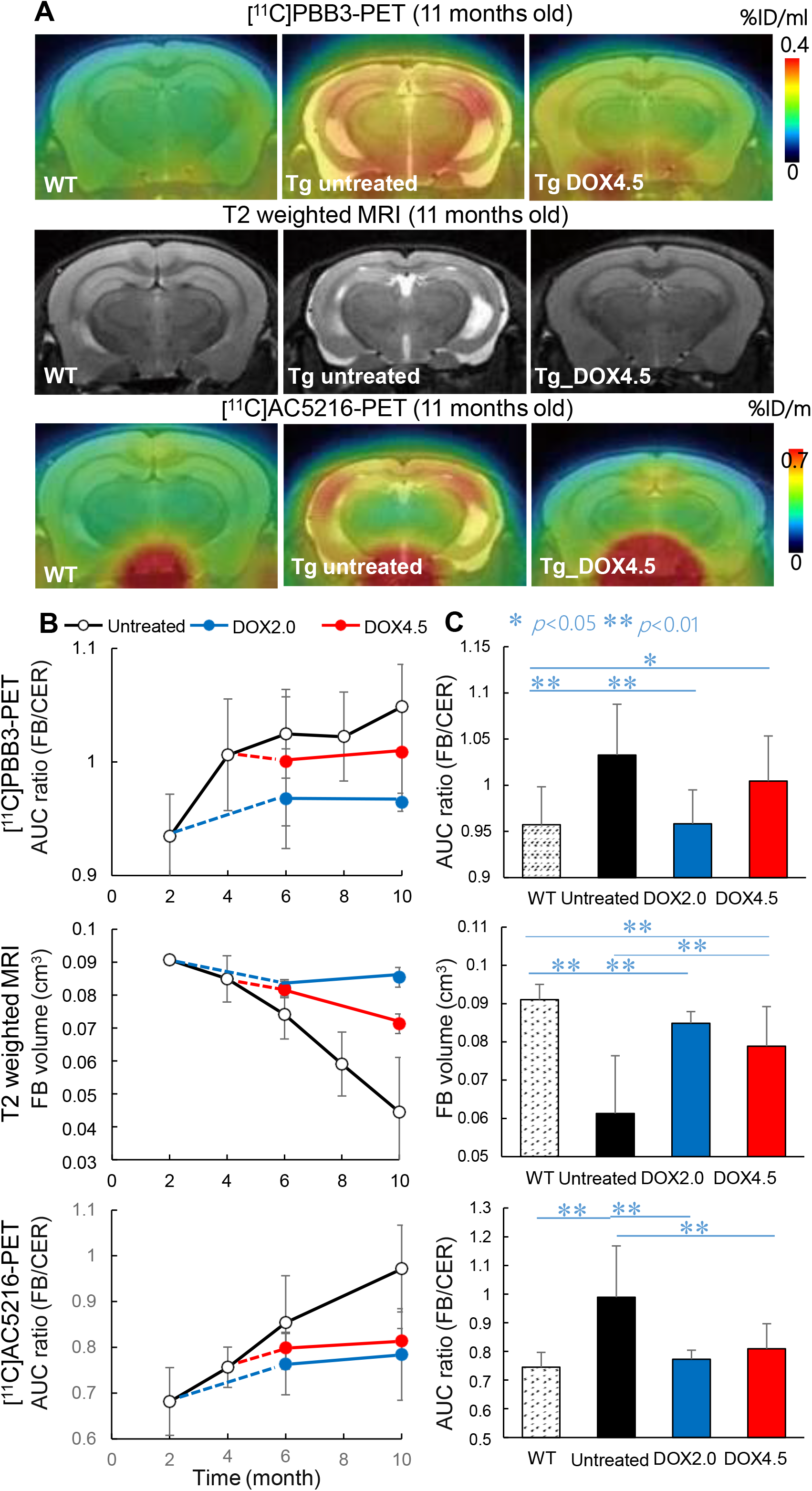
In vivo macroscopic imaging of rTg4510 (Tg) and wild-type (WT) mice at different ages. A) Representative transaxial brain images showing tau depositions by PET with [^11^C]PBB3 (top row), structural changes by volumetric T2-weighted MRI (middle row), and TSPO-positive inflammatory gliosis by PET with [^11^C]AC5216 (bottom row) in untreated WT and Tg mice and Tg mouse treated with doxycycline from 4.5 months of age (DOX4.5). PET images were generated by averaging radioactivity at 30 – 60 min after radioligand injection, and were superimposed on volumetric MRI data. The radioligand retention was expressed as percentage of injected dose per unit tissue volume (%ID/ml). B) Time-course changes in binding of [^11^C]PBB3 (top panel) and [^11^C]AC5216 (bottom panel) in the forebrain (FB; a combined neocortex and hippocampal region) estimated as a ratio of areas under the curve (AUCs) between the FB and cerebellum (CB) in time-radioactivity plots, and FB volumes (middle panel) in Tg mice, which were untreated (black symbols) and treated with doxycycline from 2.0 (DOX2.0; blue symbols) and 4.5 (DOX4.5; red symbols) months of age. C) FB-to-CB AUC ratios for [^11^C]PBB3 (top panel) and [^11^C]AC5216 (bottom panel), and FB volumes (middle panel) in wild-type, and untreated and treated Tg mice. Error bars represent S. D.

To examine the mechanistic links of tau fibrillogenesis to local inflammation and shrinkage of the brain, rTg4510 mice were treated with doxycycline, a blocker of the binding of an inducible transcriptional activator to the transgene promoter, resulting in almost complete suppression of the transgenic tau expression ^13^. Initiation of the treatment at 2 months of age profoundly halted the progression of tau and inflammatory pathologies and atrophy (Figure 1), supporting causal relationships among these alterations. Treatment commenced at 4.5 months of age also markedly attenuated the induction of neuroinflammation and reduction of regional volumes, but did not significantly repress tau accumulations (Figure 1), raising the possibility that the transgene inactivation protected tangle-bearing neurons against death. Meanwhile, it remained to be further investigated whether soluble intermediates of tau assemblies or detrimental microgliosis could provoke loss of these neurons.

### Rapid turnover of tau fibrils as a consequence of tangle generation in parallel with loss of tangle-burdened neurons

We subsequently conducted intravital two-photon laser fluorescence microscopy of the rTg4510 mouse neocortex to monitor the same individual tau inclusions over several weeks. Neuronal tau deposits were captured by intravenously administered unlabeled PBB3 as documented elsewhere ^12^, and in-vivo labeling of blood vessels with sulforhodamine 101 (SR101) provided ‘street addresses’ of the PBB3-illuminated tau tangles for a longitudinal analysis (Supplemental Figure 1). Weekly imaging sessions for mice from 6.5 to 7.5 months of age revealed the appearance and disappearance of the inclusions with these two events often occurring serially in a single cell only a few weeks apart (Figure 2 A, B and Supplemental Figure 2), indicating a rather prompt loss of neurons after the emergence of PBB3-positive tau aggregates. Notably, approximately 50% of newly generated tangles disappeared within 2 weeks, and this loss was complemented by even newer tau aggregates, leading to apparently stable net counts of tau deposits during the observation period (Figure 2B, C).

**Figure 2.**
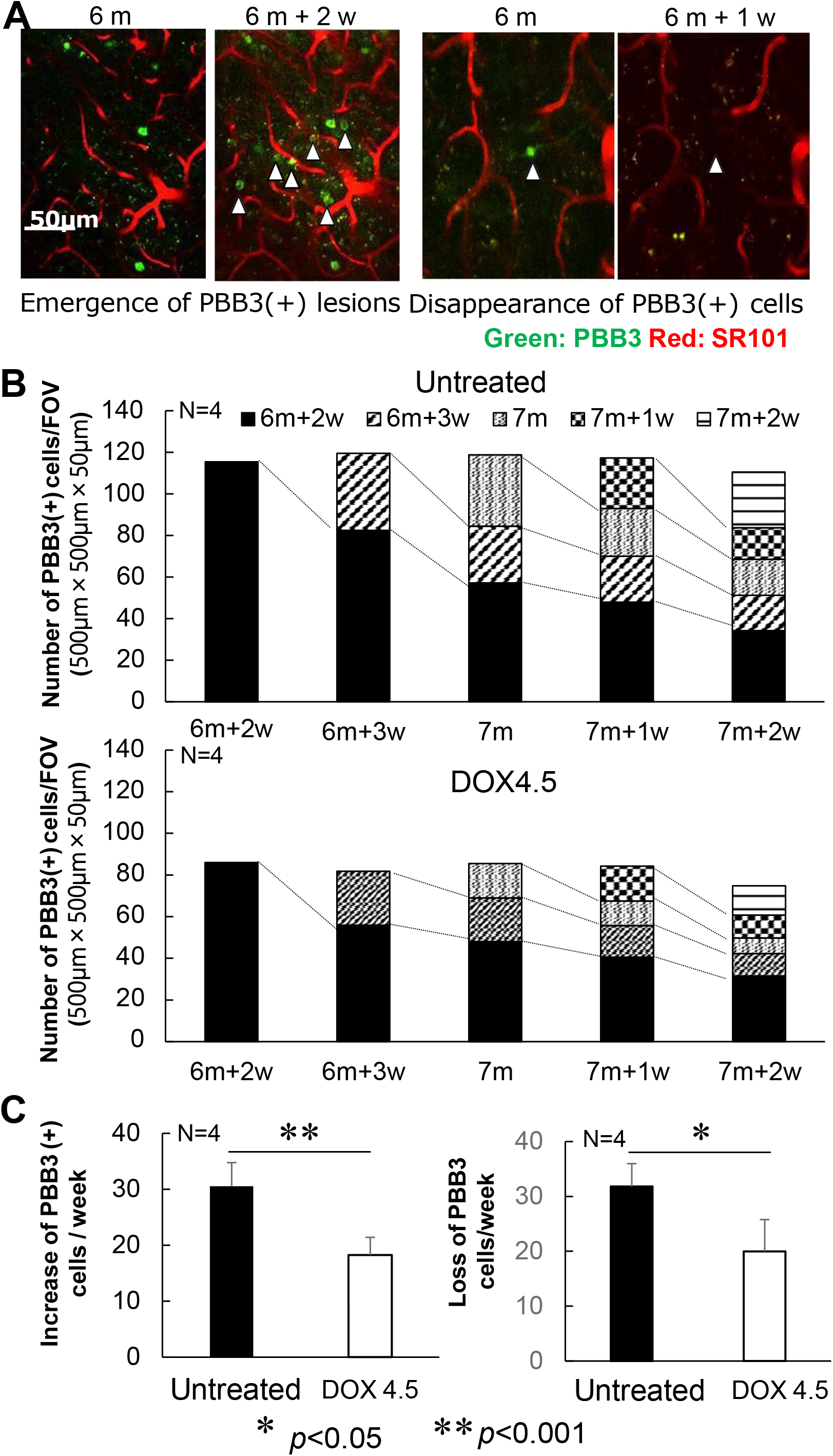
Time-course analyses of the tau fibril dynamics in rTg4510 (Tg) mouse brains by in-vivo two-photon laser fluorescence microscopy. A) Longitudinal imaging of individual tau lesions labeled with intravenously injected PBB3 (green) along with SR101-labeled blood vessels (red) at the cortical surface from 6 months of age. Left and right composites of two panels display emergence and disappearance of PBB3-positive tau lesions (arrowheads), respectively, during the observation period. B) Chronological changes in the number of PBB3-positive cells per field of view (FOV) in untreated Tg mice (top panel) and Tg mice treated with doxycycline from 4.5 months of age (DOX4.5; bottom panel). Solid columns in the stacks represent the number of tau lesions existing from 6 months of age, and columns with the same non-solid texture show chronological changes in the number of tau lesions emerging at the following week intervals. The data are averages of four mice in each treatment group. C) Rates of emergence (left) and loss (right) of PBB3-positive lesions estimated as the number of events per FOV per week. Solid and open columns indicate data obtained from Tg mice in untreated and DOX4.5 groups, respectively. Error bars represent S. D. *p < 0.005 by t-test.

We then initiated treatment of rTg4510 mice at 4.5 months of age with doxycycline, and monitored tau fibrillogenesis weekly from 6.5 to 7.5 months of age. Again, turnover of tau fibrils was observed as a result of new generation and the following disappearance of tangles, and the transgene inactivation led to a modest decrease in the net counts of tau deposits, by ~25% relative to untreated controls (Figure 2B). However, quantitative analyses of these longitudinal data illustrated more pronounced decreases in weekly rates of new tangle productions and loss of tangles, by ~40% as compared with controls (Figure 2C). Hence, the efficacies of experimental therapeutics might be underestimated in the measurements of net PBB3 positivity by PET and optical imaging, whereas separate determinations of tangle appearance and disappearance rates provide more sensitive indices for the therapeutic modifications of tau pathologies.

### High lethality of neurons bearing a package of dense tau fibrils relative to those carrying less mature tau assemblies

Since the loss of PBB3-positive tau lesions may reflect the death of neurons with densely packed tau fibrils, it is conceivable that toxic insults take place in these neurons despite isolation of tau aggregates in a subcellular compartment. To compare the propensity of neurons with different stages of tau assemblies to pathological death, we utilized PBB3 and its derivative, PM-PBB3 ^18^, which is capable of detecting less mature tau packages than PBB3, for in-vivo fluorescence microscopy of rTg4510 mouse brains. A head-to-head comparison of real-time images yielded by systemic injections of the probes demonstrated that PM-PBB3 can illuminate many more tau lesions than PBB3 in the same field of view (Figure 3A), and PBB3-negative, PM-PBB3-positive inclusions are considered to be tau tangles at an earlier stage.

**Figure 3.**
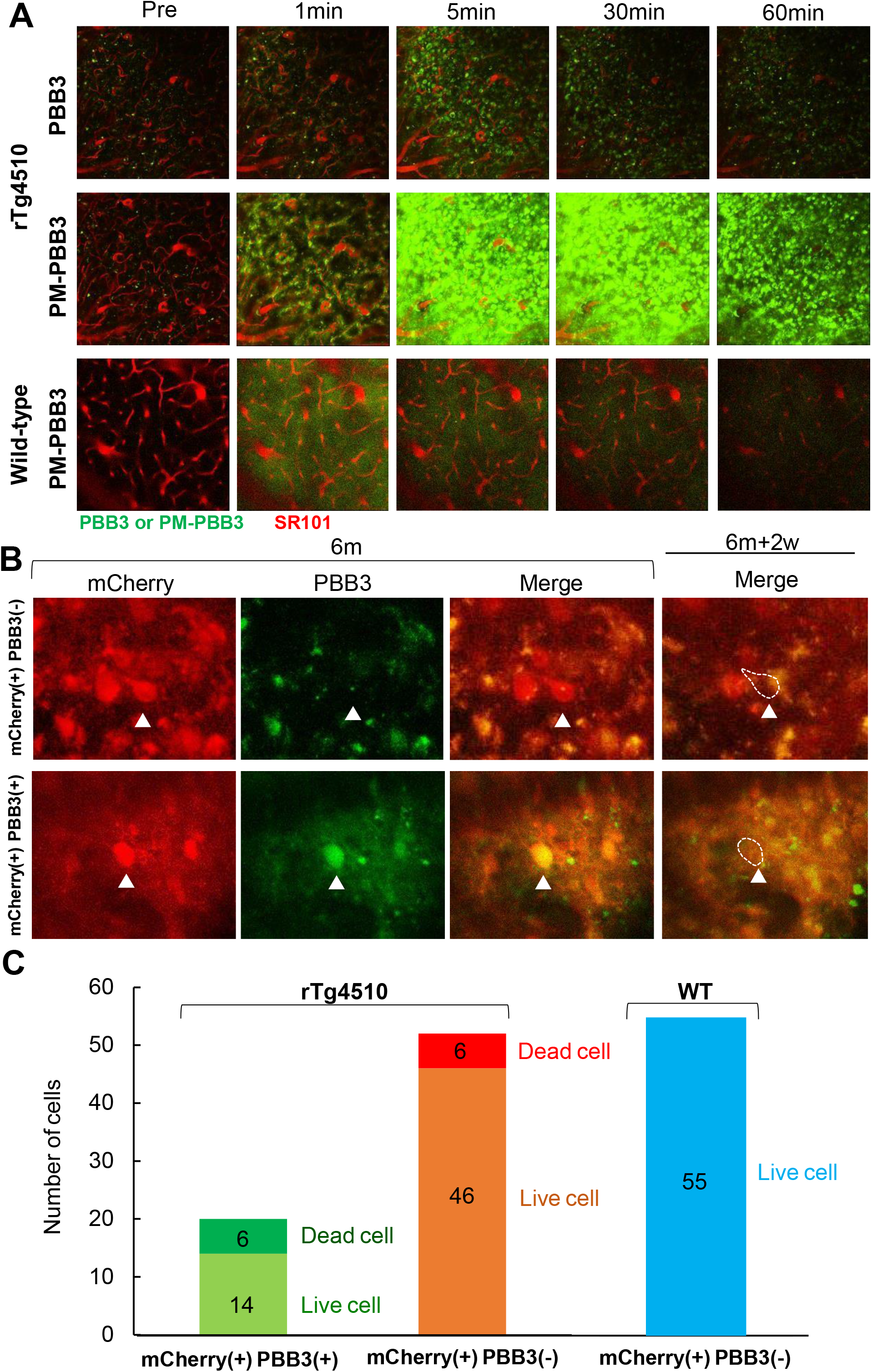
Loss of neurons bearing tau aggregates with different packing densities in the neocortex of rTg4510 mice. A) Comparison of tau lesions labeled with intravenously administered PBB3 (top panels) and PM-PBB3 (middle panels) of the same field of view of the brain of an rTg4510 mouse tracked by real-time two-photon laser microscopic imaging. Corresponding images in the brain of a wild-type mouse are also displayed (bottom panels). Tau aggregates are illuminated in green by PBB3 or PM-PBB3, and blood vessels are visualized in red by SR101. B) Disappearance of mCherry-positive neurons possessing PBB3-negative (presumably PM-PBB3-positive) tau aggregates (arrowheads in top panels) and PBB3-positive tau aggregates (arrowheads in bottom panels) during the 2-week observation period from 6 months of age. Note that almost all mCherry-positive neurons were burdened with PM-PBB3-positive tau aggregates at this age (see Supplemental Figure 3). C) Rate of loss of mCherry-positive neurons labeled (left column) and unlabeled (middle column) with PBB3 versus survival of these cells in the neocortex of rTg4510 mice during the week observation period estimated as the number of events per field of view per week. 30% of PBB3-positive and 12% of PBB3-negative neurons disappeared during this period. Unlike rTg4510 mice, no loss of mCherry-positive neurons was observed in wild-type (WT) mice (right column). The numbers in each stacked column denote the numbers of live or dead cells.

In order to determine the fraction of PM-PBB3-positive cells among the total neuronal population, a fluorescence protein, mCherry, was expressed in neurons by inoculation of an adeno-associated viral (AAV) vector carrying a gene coding this protein downstream to the neuron-specific synapsin-1 promoter. In-vivo assays revealed that all mCherry-positive neurons were also labeled with PM-PBB3 in the neocortex of rTg4510 mice at 6 months of age (Supplemental Figure 3), providing a rationale for assessing mCherry signals as surrogates of PM-PBB3-detectable tau aggregates in these mice. A weekly investigation of individual neurons in living mice showed that the number of PBB3-negative neurons bearing less mature tau fibrils was 2.5-fold larger than that of PBB3-positive neurons, but a comparable number of neurons died in these two cellular cohorts at 2 weeks after the baseline observation (Figure 3B, C), suggesting high lethality of neurons enriched with PBB3-positive mature tangles in comparison with those possessing relatively immature forms of fibrillar tau deposits. Hence, there should exist a mechanism of neuronal death accompanied by the condensation and maturation of tau aggregates.

### Engulfment of tangle-burdened neurons by phagocytic microglia followed by neuronal death

Since the frequent loss of neurons burdened with densely packed tau tangles left no remnants such as extracellular tau deposits dubbed ghost tangles, all cellular components including tangles are likely to be removed by phagocytic glia. This clearing process may begin non-cell-autonomously in live neurons, or promptly after cell-autonomous death of neurons. To determine the chronology of phagocytosis and death of neurons in this circumstance, we expressed mCherry in neurons and enhanced green fluorescence protein in microglia driven by promoters of synapsin-1 and CD68, respectively, by means of intracranially inoculated AAV vectors.

In-vivo double labeling of these two cellular populations in rTg4510 mice revealed engulfment of healthy-looking neurons by hypertrophic cell bodies or processes of microglia (Figure 4A B). Intravenous administration of PBB3 or PM-PBB3 intensified fluorescence signals in these neurons, indicating the presence of tau tangles (Supplemental Figure 4). Daily follow-up observations of these alterations demonstrated that the engulfed neurons shrank and then vanished within a week (Figure 4C). Weekly monitoring also serially captured initiation of the microglial engulfment and the disappearance of a single neuron with these two events occurring approximately 2 weeks apart (Supplemental Figure 5). As a result of the phagocytic removal of neuronal somas, no tau fibrils were left detectable by the tau probes. Thus, these results represent a direct demonstration of an essential role played by microglia in tau-triggered neuronal death without formation of ghost tangles.

**Figure 4.**
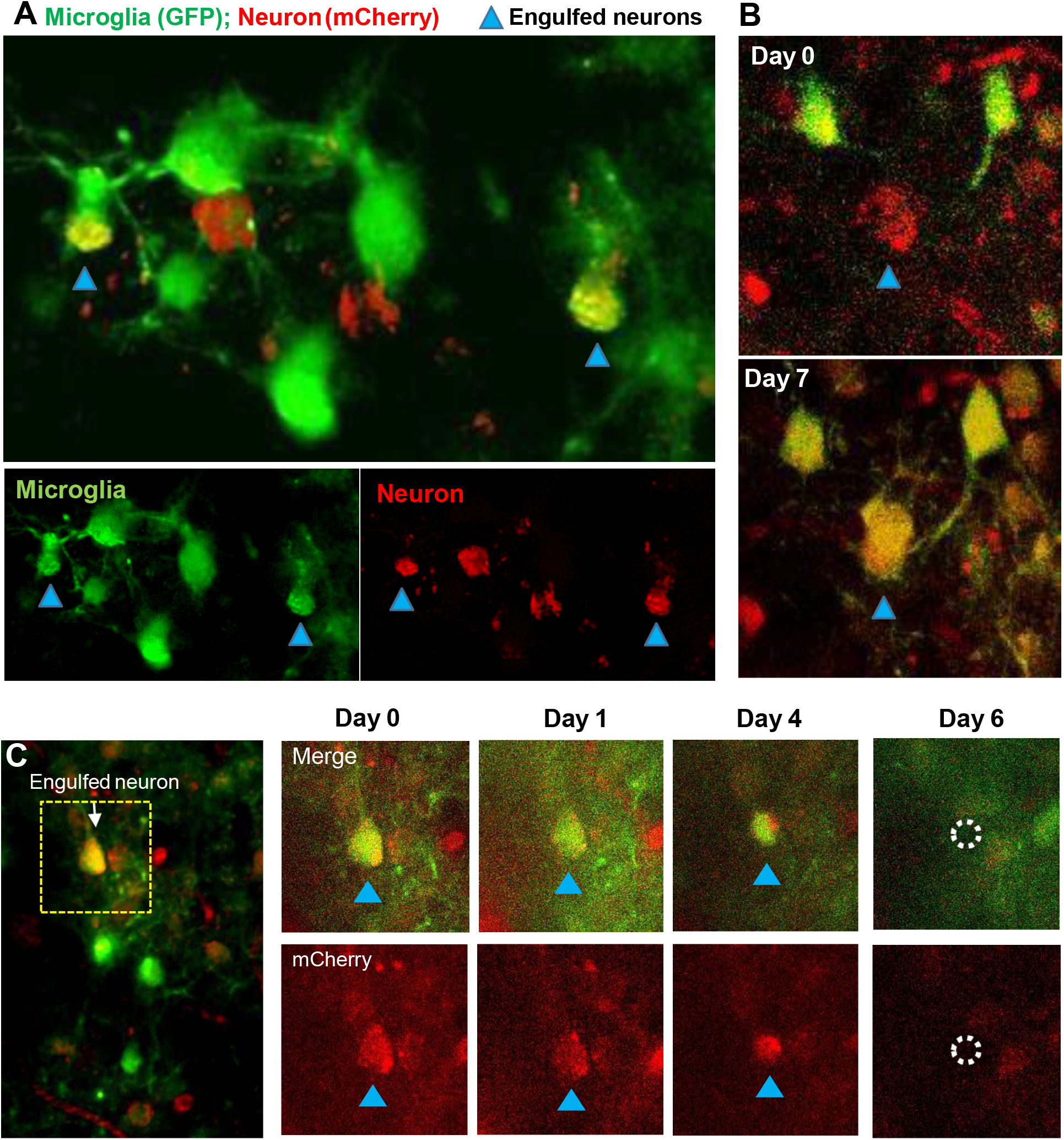
Longitudinal intravital two-photon laser microscopy demonstrating engulfment and phagocytosis of neurons by microglia in the rTg4510 mouse neocortex. A)Engulfment of mCherry-expressing neuron by hypertrophic processes of GFP-expressing microglia (arrowheads) in an rTg4510 mouse at 6 months of age. Merged two-channel (top) and separate green and red channel (bottom) images are displayed. B) Weekly monitoring of neuronal engulfment by microglia. A neuron (arrowheads) is not in close contact with microglia at baseline (top), but is engulfed by an enlarged process stretching from neighboring microglia 1 week later (bottom). Note that almost all engulfed mCherry-positive neurons are burdened with PM-PBB3-positive tau aggregates (Supplemetal Figure 4). C) Loss of a neuron after being engulfed by phagocytic microglia captured by daily microscopic imaging. A neuronal cell body engulfed by microglia at baseline (arrow in left large field of view and arrowhead in small panels) subsequently exhibited shrinkage at Day 4 and elimination at Day 6. Upper and lower panels show merged 2-channel and single red-channel fluorescence photomicrographs, respectively.

**Figure 5.**
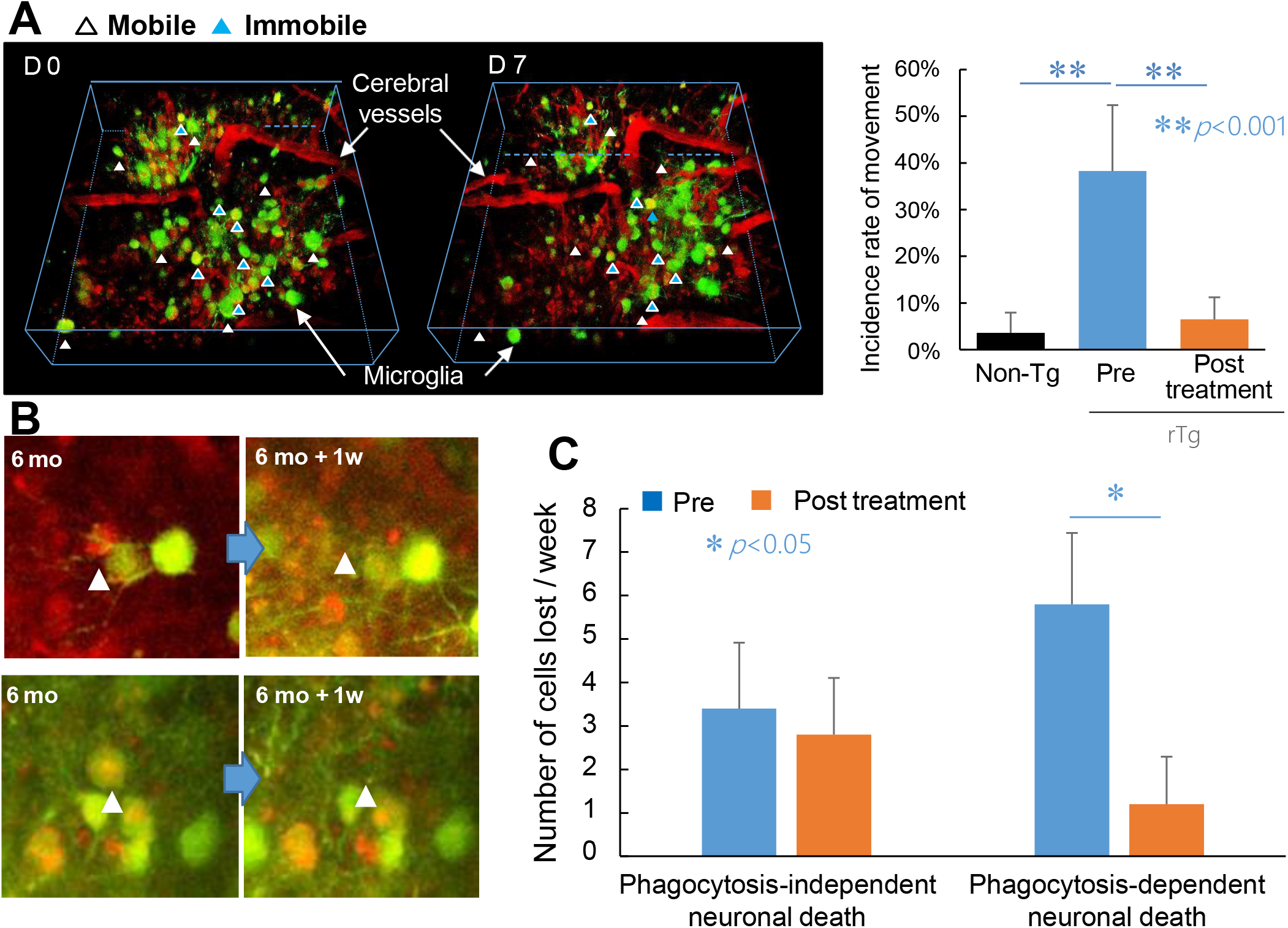
Pharmacological suppression of phagocytic microgliosis leading to survival of neurons bearing tau tangles. A) Effects of a TSPO ligand, Ro5-4864, on the in vivo mobility of microglia in the rTg4510 mouse brain. 3-D images of GFP-expressing microglia (green) and SR101-labeled blood vessels (red) were generated by Z-stacking of multiple X-Y planes (left panel). Immobile microglia located in the same position at baseline and Day 7 are indicated by blue arrowheads, and microglia with high mobility entering or leaving the 3-D field of view at Day 7 are indicated by white arrowheads. The vast majority of microglia in wild-type control mice did not display mobility, while highly mobile cells accounted for approximately 1/3 of the microglial population in rTg4510 mice at 6 months of age before treatment. This hypermobility was markedly suppressed by one-week treatment of mice with Ro5-4864. N = 5 in each group. B) Loss of mCherry-positive neurons (red) in a manner independent of (upper panels) and dependent on (lower panels) phagocytosis by GFP-positive microglia (green). A weekly two-photon microscopic assay captured disappearance of some neurons (arrowheads) at 1 week after the baseline observation without contacting microglia (upper panels), while other neurons were eliminated at 1 week after being engulfed by phagocytic microglia (lower panels). C) Rates of phagocytosis-independent and phagocytosis-dependent neuronal deaths estimated as number of events per field of view per week during 1-week observations before (blue columns) and after (orange columns) initiation of the treatment with Ro5-4864. *p < 0.05 by t-test. Error bars represent S. D.

### Pharmacological intervention in deleterious microgliosis leading to suppression of phagocytosis-dependent loss of neurons

We further employed the intravital microscopic imaging system to examine the notion that neurons are able to survive if they are prevented from being phagocytosed by aggressive microglia. As shown in the current PET experiments, together with previous works ^6,16,17^, a notable feature of the deleterious microglia is a high-level expression of TSPO. In light of the fact that TSPO is a component residing in the mitochondrial outer membrane responsible for the incorporation of cholesterol in the mitochondrial steroidogenesis ^19,20^, TSPO could act against aggressive microglial phenotypes via immunosuppressive steroids, although this mechanism in natural history seems to be insufficient for arresting microglia overdriven by misfolded tau species.

Driven by this hypothetical view, rTg4510 mice were treated with an agonistic TSPO ligand, Ro5-4864, which has been reported to enhance microglial production of steroids ^21,22^. For Ro5-4864-treated and untreated mice, a longitudinal microscopic comparison of microglial locations was performed between baseline and 1 week post-treatment. Assays of untreated animals showed that microglia are barely mobile in the neocortex of non-transgenic mice at 6 months of age, in sharp contrast with microglia with high mobility accounting for ~40% of the total microglial population in rTg4510 mice at the same age (Figure 5A). Significantly, this hypermobility of microglia in rTg4510 mice was drastically inhibited by 1-week treatment with Ro5-4864 (Figure 5A). We also conducted in-vivo double-labeling of neurons and microglia to quantify the numbers of neurons dying in manners dependent on and independent of microglial phagocytosis within 1 week (Figure 5B). It is noteworthy that the majority of neuronal death was phagocytosis-dependent in untreated rTg4510 mice (Figure 5C), and this was markedly blocked by the Ro5-4864 treatment, in contrast to the absence of overt effects of this agent on non-phagocytic neuronal loss (Figure 5C). According to these findings, a large proportion of tangle-bearing neurons can survive if microglial attacking of these cells is therapeutically deactivated.

### Tagging of neuronal somas with complement MFG-E8 and C1q as a putative trigger of immediate phagocytosis

The present demonstration of phagocytic removal of neurons carrying tau lesions has highlighted the possibility that vital neurons could be mislabeled with molecular tags, which guide aggressive microglia toward the lethal engulfment of neurons. In line with this postulation, we found abundant immunolabeling of the complements C3 and C1q chiefly in somas and synapses, respectively, of neurons in the brains of rTg4510 mice at 6 months of age, in clear distinction from the faint immunoreactivity in non-transgenic controls (Figure 6). The distribution of these complements was confined to the forebrain, in agreement with the localization of tau depositions in these animals ^13–15^.

**Figure 6.**
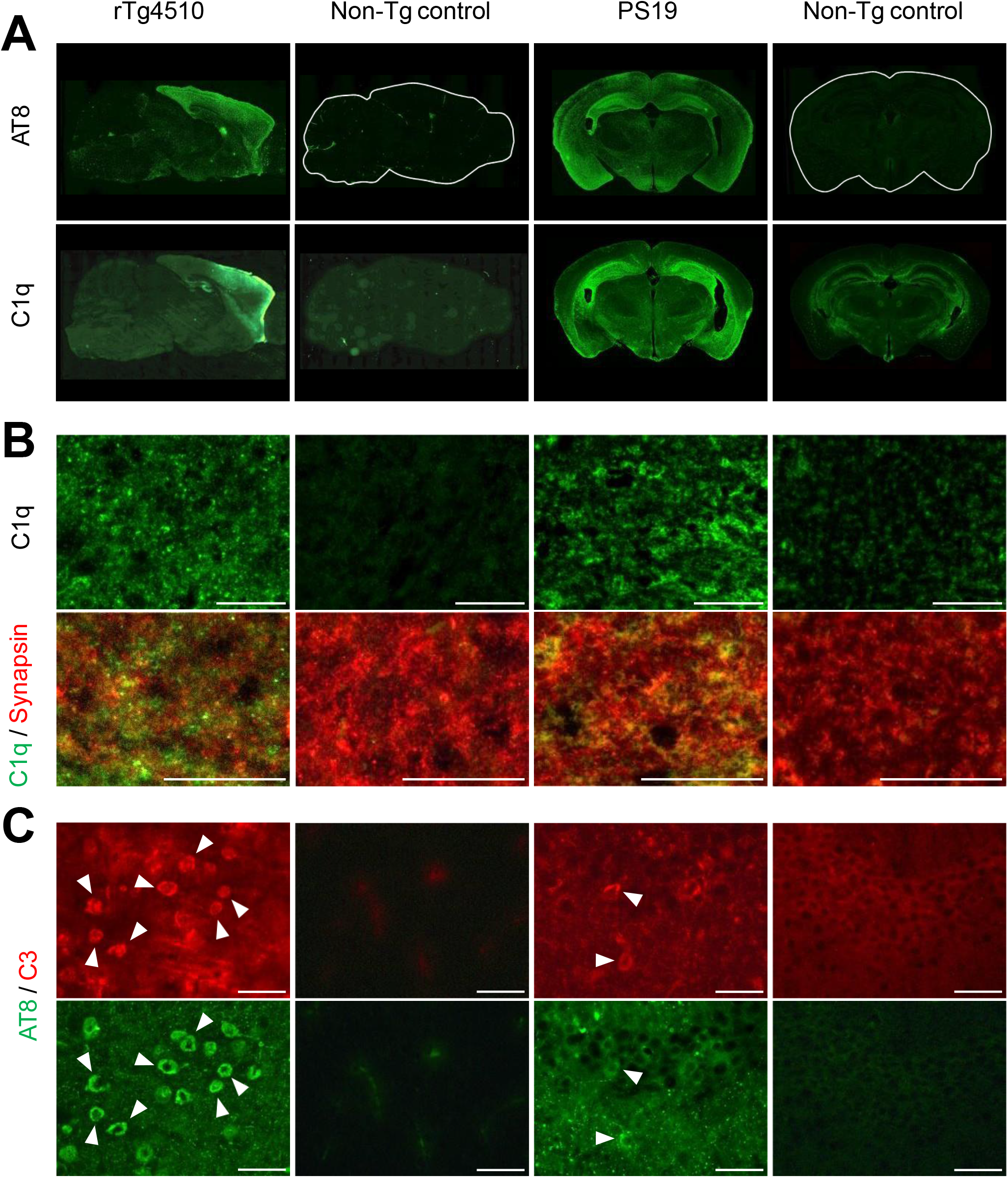
Accumulation of complements, C3 and C1q, in distinct subcellular compartments of neurons in the brains of rTg4510 and PS19 mice. A) Low-power photomicrographs showing immunolabeling of phosphorylated tau (AT8, top) and C1q (bottom) in sagittal sections of rTg4510 (left) versus age-matched non-Tg control (left middle) mouse brains and coronal sections of PS19 (right middle) versus age-matched non-Tg control (right) mouse brains. B) High-power images showing immunolabeling of C1q (top) and double immunolabeling of C1q (green in bottom) and synapsin (red in bottom) in the neocortex of rTg4510 (left) versus age-matched non-Tg control (left middle) mice and oriens layer of the hippocampus of PS19 (right middle) and age-matched non-Tg control (right) mice. C) High-power images of double immunolabeling of C3 (top) and phosphorylated tau (AT8, bottom) in the neocortex of rTg4510 (left) versus age-matched non-Tg control (left middle) mice and oriens layer of the hippocampus of PS19 (right middle) and age-matched non-Tg control (right) mice. Arrowheads indicate neuronal somas doubly immunolabeled with AT8 and anti-C3 antibody. Scale bars, 50 μm (B, C).

To examine whether this pathological decoration of neuronal components with complements commonly occurs in different tauopathy models, we also analyzed immunostaining of these complements in the brains of another tau transgenic strain, PS19, which overexpresses human tau proteins with the P301S FTDP-17-*MAPT* mutation ^6,12,14^. Somatic C3 and synaptic C1q staining was noted in neurons localized to the hippocampus and entorhinal cortex of PS19 mice, regionally overlapping with fibrillary tau pathology, microgliosis and loss of neurons (Figure 6). Furthermore, diffuse, scattered C1q immunoreactivity and C3 immunolabeling of cell bodies were noted in the hippocampus of patients with AD and senile dementia of neurofibrillary tangle type, which is characterized by accumulations of tau fibrils but lacks marked amyloid-β (Aβ) deposits (Supplemental Figure 6).

These findings implied the involvement of complements in recognition of tangle-burdened neurons by microglia, but the labeling of numerous neuronal somas with C3 raised a possibility that this decoration may not promptly provoke phagocytic elimination of the labeled neurons by microglia. To identify molecular tags immediately inducing phagocytic removal of affected neurons, we treated rTg4510 mice with an inhibitor of colony-stimulating factor 1 receptor (CSF1R), PLX-3397 (PLX). Oral administration of PLX (290 mg/kg chow) to the rTg4510 mice at the age of 4 – 6 or 7 - 8 months resulted in a partial depletion of microglia (Supplemental Figure 7A) and suppression of neurons as assessed by volumetric MRI (Figure 7A) and postmortem immunohistochemistry (Supplemental Figure 7B), indicating that this dosage of PLX is capable of diminishing primary phagocytosis of neurons loaded with tau fibrils.

**Figure 7.**
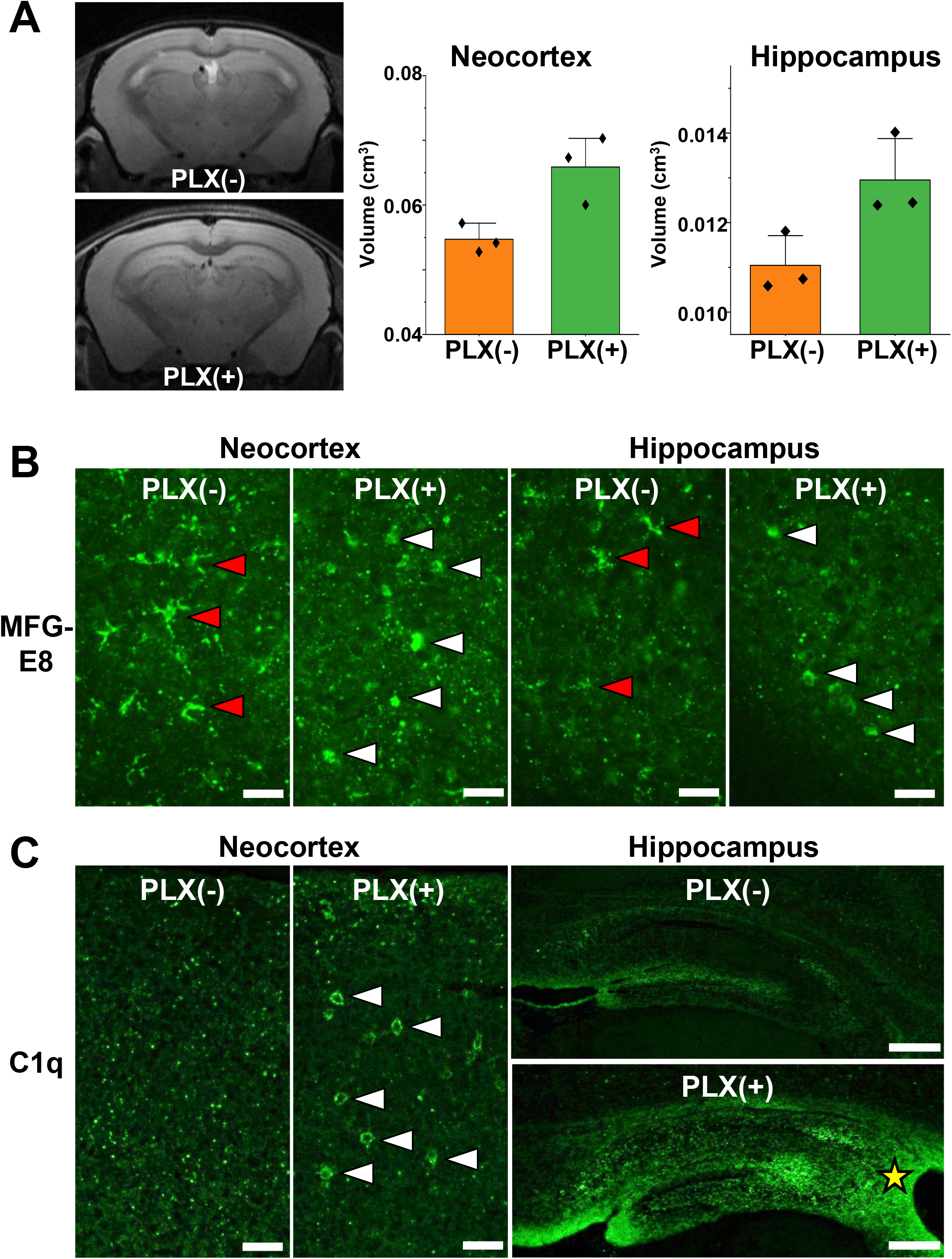
Suppression of neuronal death in concurrence with increased labeling of neuronal somas with MFG-E8 and C1q, indicating sparing of opsonin-tagged neurons from phagocytic elimination. A) Coronal T2-weighted MR images of the 6-month-old rTg4510 mouse brains, containing the neocortex, hippocampus, and thalamus (left) and quantification of neocortical and hippocampal volumes (right; n = 3 in each treatment group). The MRI data were acquired in an untreated condition [PLX(-)] or following PLX treatment over 2 months [PLX(+)]. There were main effects of treatment [F(1, 8) = 14.90, p = 0.005] and region [F(1, 8) = 811.7, p < 0.0001] and interaction between these two factors [F(1, 8) = 7.485, p = 0.03] as examined by 2-way ANOVA. Filled rhombuses denote individual values, and a vertical line above each column indicates SD. Immunofluorescence staining of MFG-E8 in brain sections derived from rTg4510 mice. Brain tissues were obtained from 8-month-old mice without treatment [PLX(-)] or following PLX treatment over 1 month [PLX(+)]. MFG-E8 immunoreactivity was localized to putative microglia (red arrowheads) in the untreated condition, while the labeling was markedly decreased in microglia and increased in neuronal somas (white arrowheads) by the PLX treatment. C) Immunofluorescence staining C1q in brain sections sub-adjacent to those displayed in B. The labeling was primarily localized to synapses as punctate signals in the untreated condition (see also Figure 6B) and was profoundly increased in neuronal somas (white arrowheads) by the PLX treatment in the neocortex. A marked enhancement of C1q labeling by the treatment was also observed in fiber bundles surrounding the hippocampus, including the fimbria (yellow star) nearby the lateral ventricle. Scale bars, 50 μm (B and left and middle panels in C) and 250 μm (right panels in C).

We then immunohistochemically analyzed molecules potentially acting on neurons as ‘eat-me’ signals. Among these components, milk fat globule-epidermal growth factor 8 protein (MFG-E8) was found to decorate neurons for a rapid engulfment by microglia. In the neocortex and hippocampus of PLX-untreated rTg4510 mice, MFG-E8 immunostaining was mostly observed in microglia, whereas labeling of neuronal cell bodies with MFG-E8 was barely observed (Supplemental Figure 8). By contrast, PLX treatment profoundly reduced MFG-E8-positive microglia and increased MFG-E8-tagged neuronal somas (Figure 7B). Hence, microglia remaining in the PLX treatment could produce and release MFG-E8 but was incapable of phagocytosing MFG-E8-labeled neurons. This finding also indicates that MFG-E8 attached to cell bodies of tangle-burdened neurons acts as an ‘eat-me’ signal to rapidly give rise to microglia-mediated removal of these neurons, impeding the detection of MFG-E8-labeled neuronal soma in the untreated condition.

We also compared the labeling of neurons with complements between untreated and PLX-treated conditions. While C1q was mostly localized to synaptic compartments with few somatic labeling in the untreated mouse neocortex and hippocampus, neuronal cell bodies with the attachment of this complement were markedly increased by the PLX treatment (Figure 7C). Similar to MFG-E8, C1q may accordingly mediate the loss of neurons as an immediate inducer of the primary phagocytosis of neuronal somas. Unlike C1q, C3 immunolabeling was frequently detectable in neuronal somas of the untreated rTg4510 mouse forebrain and was further increased following the PLX treatment, in line with the rescue of these neurons (data not shown). Therefore, C3 on the neuronal cell bodies is unlikely to promote a quick attraction of phagocytic microgliosis.

C1q, an initiator of the classical complement cascade, is known to label unnecessary synapses to attract phagocytic microglia in the development-related pruning of excess neural connections ^23,24^. Recently, C1q and microglia have also been found to mediate phagocytic loss of synapses encompassing Aβ plaques in a mouse model ^25^. Moreover, MFG-E8 is a factor binding to phosphatidylserine presented on the surface of apoptotic cells. A recent report documented that non-apoptotic neuronal death triggered by tau depositions could be accompanied by the MFG-E8 labeling in a transgenic mouse model of tauopathies ^26^. We accordingly conceive that aberrant tagging of neurons with MFG-E8 and C1q direct labeled neurons to prompt ‘eating’ by activated microglia.

## DISCUSSION

The present work has delineated spatiotemporal and causal relationships among tau fibrillogenesis, inflammatory microgliosis, and neuronal death by multimodal macroscopic and microscopic imaging assessments of these pathological processes in a living tauopathy model. Importantly, the rapid turnover of tau fibrils is present in the brain of this model and is attributed to a relatively short life of tau tangles after becoming detectable by the imaging agent. The disappearance of tau tangles reflects the loss of tangle-burdened neurons, which has been found to be primarily mediated by phagocytic microglia. This phagocytosis is likely to cause the death of viable neurons, according to the rescue of tangle-bearing neurons by pharmacological suppression of microglial mobility and phagocytic activity. Notably, microglia remaining after the PLX treatment was devoid of the ability to phagocytose neurons but was still capable of releasing complements and related immune signals. In this condition, a relatively small subset of neuronal somas loaded with tau tangles was found to be labeled with MFG-E8 and C1q. This tagging was barely observed in the untreated mice, indicating that MFG-E8 and C1q exposed on cell bodies of neurons rapidly induce vanishment of these neurons via opsonization by phagocytic microglia.

A previous two-photon laser microscopic assay of rTg4510 mice focused on apoptotic signaling triggered by neuronal tau depositions and illustrated that formation of thioflavin-S-positive tau tangles occurs promptly after caspase activation and may lead to attenuation of the caspase actions ^27^. This change prevents neurons from apoptotic cell death during an observation period of 2 – 5 days ^27^, suggesting that loss of tangle-burdened neurons caused by apoptotic processes could be a rare pathological event. In the current work, however, monitoring of individual neurons and tau lesions over a longer term revealed that approximately 50% of neurons were eliminated within 2 weeks after the emergence of dense tau aggregates, which is underlain by non-apoptotic, non-cell-autonomous mechanisms critically involving phagocytic microglia.

The PLX treatment led to the revelation that opsonins hardly detectable on neurons in the untreated condition should act as ‘eat-me’ signals prompting phagocytic removal of these neurons in a short period. The phagocyte-dependent neuronal cell death has been characterized by ‘eat-me’ signals exposing phosphatidylserine on the neuronal cell surface ^28^, and the release of MFG-E8 and its binding to phosphatidylserine facilitates engulfment of live neurons ^29^. In a tauopathy mouse model expressing P301S tau driven by the Thy-1 promoter ^30^, living neurons with tau inclusions highly exposed phosphatidylserine and MFG-E8 expression was elevated in the brains ^26^. Furthermore, inhibition of MFG-E8 binding to the vitronectin receptor subunit abrogated phagocytosis and rescued living neurons ^26^. The current data have provided more compelling evidence for the critical involvement of this molecule in the rapid engulfment of neuronal somas affected by tau pathologies. Likewise, C1q attached to neuronal somas was identified as a potent inducer of opsonization, leading the neurons to immediate death. It is also noteworthy that C1q labeling was intensified on the nerve fibers in several hippocampal subregions, including the fimbria in the vicinity of the lateral ventricle, in the PLX-treated versus untreated condition (Figure 7C), which implies that prompt loss of these fiber bundles may accelerate the ventricular enlargement in the untreated rTg4510 mice.

In the absence of the PLX treatment, neuronal somas and synapses were frequently labeled with C3 and C1q, respectively, supporting the possibility that these tags may persist for a long term without driving aggressive microglial responses. Accumulations of C1q and C3 have been noted as common pathological features of rTg4510 and PS19 mice and human tauopathies, indicating complement-driven deteriorations of living neurons as a consensus mechanism of neurodegenerative changes in these strains and species. Indeed, upregulation of C3 and its receptor, C3aR, has recently been reported in the brains of AD and non-AD tauopathy patients and PS19 mice, and disruptions of the C3 signaling pathway have led to attenuation of neuroinflammation and neurodegeneration of PS19 mice ^31^. In addition, C1q has been observed to tag synapses affected by tau pathologies in PS19 mice, presumably provoking removal of these synapses by microglia ^32^. Somatic C3 and synaptic C1q labeling, however, may lead to mild and slow aggravation of the neuronal integrity relative to decorations of neuronal somas with MFG-E8 and C1q. Importantly, these findings and their indications bring a view that different neuronal compartments may be attacked by glia with specific phenotypes, which is mediated by unique opsonins. Such associations are exemplified by the notion that a neuronal soma tagged with MFG-E8 and C1q could be rapidly targeted by a microglial cell exerting high phagocytic activity, in contrast to less aggressive microglia responding to C3-labeled somas. A recent study demonstrated that specific glial types are responsible for phagocytosing subcellular compartments of a dead neuron in a non-overlapping manner ^33^ and the present data expand this view to the opsonin-mediated mechanism underlying the primary phagocytosis of living neurons.

The molecular and cellular elements linking the tau deposition to the opsonization and activation of phagocytic microglia remain to be identified, but it can be postulated that the accumulation of C1q is induced by a mechanism resembling synaptic pruning during postnatal development ^23,24^. Soluble tau species may deteriorate postsynaptic integrity as documented previously ^34,35^, and resultant aggravation of synaptic activities could trigger tagging of these subneuronal compartments with C1q produced by either astrocytes or microglia ^36,37^. As phagocytic microglia were shown to target neurons with impaired activities ^37^, the preferential elimination of neurons burdened with PBB3-positive tau tangles by microglia in the rTg4510 neocortex may indicate functional deficits of these neurons. Indeed, our pilot ex-vivo NanoSIMS (secondary ion mass spectrometry) analysis coupled with transmission electron microscopy demonstrated that the utilization of systemically administered isotope-labeled glucose is disturbed in neuronal somas possessing putative tangles with high maturity (Supplemental Figure 7). Therefore, tangle-bearing neurons could be viable but may not be metabolically intact, leading to the selection of these cells for the targeted phagocytosis via opsonization with MFG-E8 and C1q.

It should also be noted that the formation of PBB3-detectable tau tangles in the hippocampus and neocortex of PS19 mice is much less frequent than those seen in rTg4510 mice ^12,14^. This could imply that the tangles in PS19 mice have a very short lifespan due to the elimination of neurons immediately after the maturation of tau lesions into a PBB3-positive stage and tagging of these tangle-loaded cells with MFG-E8 and C1q. Another possibility is that immature tau aggregates act as strong inducers of microglia-mediated neuronal loss in the PS19 model. As neurodegenerative tau pathologies in PS19 mice are mainly observed in the hippocampus, the vulnerability of hippocampal neurons to tau-triggered toxic insults may differ from that of neocortical neurons analyzed in the present microscopic imaging of rTg4510 mice. Intravital two-photon laser microscopy of the hippocampus through a cranial window is not technically available at this time, but the development of a new infrared laser system with high power and optimized wavelength and pulse width for deep-brain imaging is underway.

The previous and current evidence for the deposition of complements ^31,32,38^ and TSPO-positive microgliosis ^16^ in close association with fibrillary tau pathologies in AD and non-AD tauopathy brains supports the occurrence of phagocytosis-dependent neuronal death similar to murine models in human diseases. To demonstrate the contribution of phagocytic microglia to the loss of tau-bearing neurons in human subjects, PET imaging of tau deposition and neuroinflammation and MRI assessment of brain atrophy will be required along the course of pharmacological interventions in tau pathogenesis with a modifier of microglial phenotypes as exemplified by a TSPO ligand. A clinical study of this design would also provide proof of the mechanism of a potential drug counteracting the tau-provoked neurodegenerative cascade. For non-clinical evaluation of such a candidate, a therapeutic agent could be initially applied with the present microscopic imaging of rTg4510 mice because this assaying system allows determining rates of the generation of new tangles and disappearance of tangle-carrying neurons as well as microglial mobility and phagocytotic activity within 1 week. The long-term efficacy of the candidate would subsequently be monitored in different model mice by multi-tracer PET and MRI over several months. Based on the validity in these animals, the test drug would be applicable to a clinical study or trial with macroscopic imaging probes, including PBB3, PM-PBB3, and Ac5216, which are commonly usable for non-clinical and clinical assays ^12,18,39^.

Our results highlight the probable benefits of treatments aimed at C1q, C3, MFG-E8, and TSPO to disrupt the phagocyte-dependent clearance of neurons. Among these agents, TSPO ligands could be the closest to clinical application because some TSPO-binding compounds have been used in humans as PET probes ^39^ or a candidate anxiolytic ^40^. A recent report has documented that STAT3 is a downstream element of C3-C3aR signaling and is upregulated in PS19 mouse brains ^31^, and transcription of the TSPO gene is known to be positively regulated by STAT3 ^41^. TSPO modulates a broad range of cellular functions, and several ligands for TSPO may enhance neuroprotective versus pro-inflammatory phenotypes of microglia. Inhibition of detrimental microgliosis with a TSPO ligand would also enhance the uptake of disseminating pathological tau molecules by beneficial glial cell types. A potentially more effective approach to the elimination of neurons by aggressive microglia would be the blockade of signaling pathways mediated by MFG-E8 and C1q. Receptors of these opsonins on the microglial surface, such as integrin αvβ3 and C1q collagen-like domain binding calreticulin, could be pharmacologically blocked by small-molecule compounds ^42^, although these inhibitions may also influence other adhesive functions of diverse cell types. Moreover, the present results have demonstrated that suppression of CSF1R inducing partial depletion of microglia might preferentially deactivate or depopulate microglia with high phagocytic activities while preserving other microglial phenotypes. This strategy will require a dose optimization of the CSF1R inhibitor to target deleterious microgliosis selectively, and a PET probe for CSF1R would serve the in vivo quantification of the receptor occupancy a therapeutic ligand ^43^. Neurons loaded with tau tangles could be rescued by intervening in the CSF1R signaling, as illustrated by the PLX treatment in this study, whereas the functional integrity of the surviving neurons needs to be investigated by microscopic and macroscopic imaging modalities.

On the basis of the present findings and their indications, we propose the primary phagocytosis of an otherwise viable neuron triggered by tau depositions and subsequent opsonization mechanisms as a neurodegenerative etiology. This critical event embodied in the dialogue between tangle-laden neurons and reactive microglia is trackable by a multimodal, multiscale imaging platform, contributing to the discoveries of neuronal and non-neuronal therapeutic target molecules and pharmacological agents acting on such targets towards regulatory modifications of the neuronal lethality in the diseased conditions.

## ACKNOWLEDGEMENTS

NanoSIMS imaging was conducted at Advanced Characterization Nanotechnology Platform of the University of Tokyo, supported by “Nanotechnology Platform” of the Ministry of Education, Culture, Sports, Science and Technology (MEXT), Japan. We thank Dr. Miyuki Takeuchi at the Institute of Engineering Innovation for her support. This work was supported by AMED under Grant Number JP18dm0207018, JP19dm0207072 to M. Higuchi, AMED under Grant Number JP18dk0207026, JP19dk0207049 to M. Higuchi, Core Research for Evolutional Science and Technology (CREST; 16810071) from Japan, Science and Technology Agency JST CREST Grant Number JPMJCR1652 to M. Higuchi, Grant-in-Aid for Science Research on Innovation Areas (“Brain Protein Aging” 26117001 and “Singularity Biology”19H05437 to N.S.) and Scientific Research (C) (19K06896 to N.S.) from the Ministry of Education, Culture, Sports, Science and Technology, Japan.

## Legends for Supplemental Figures

**Supplemental Figure 1.**
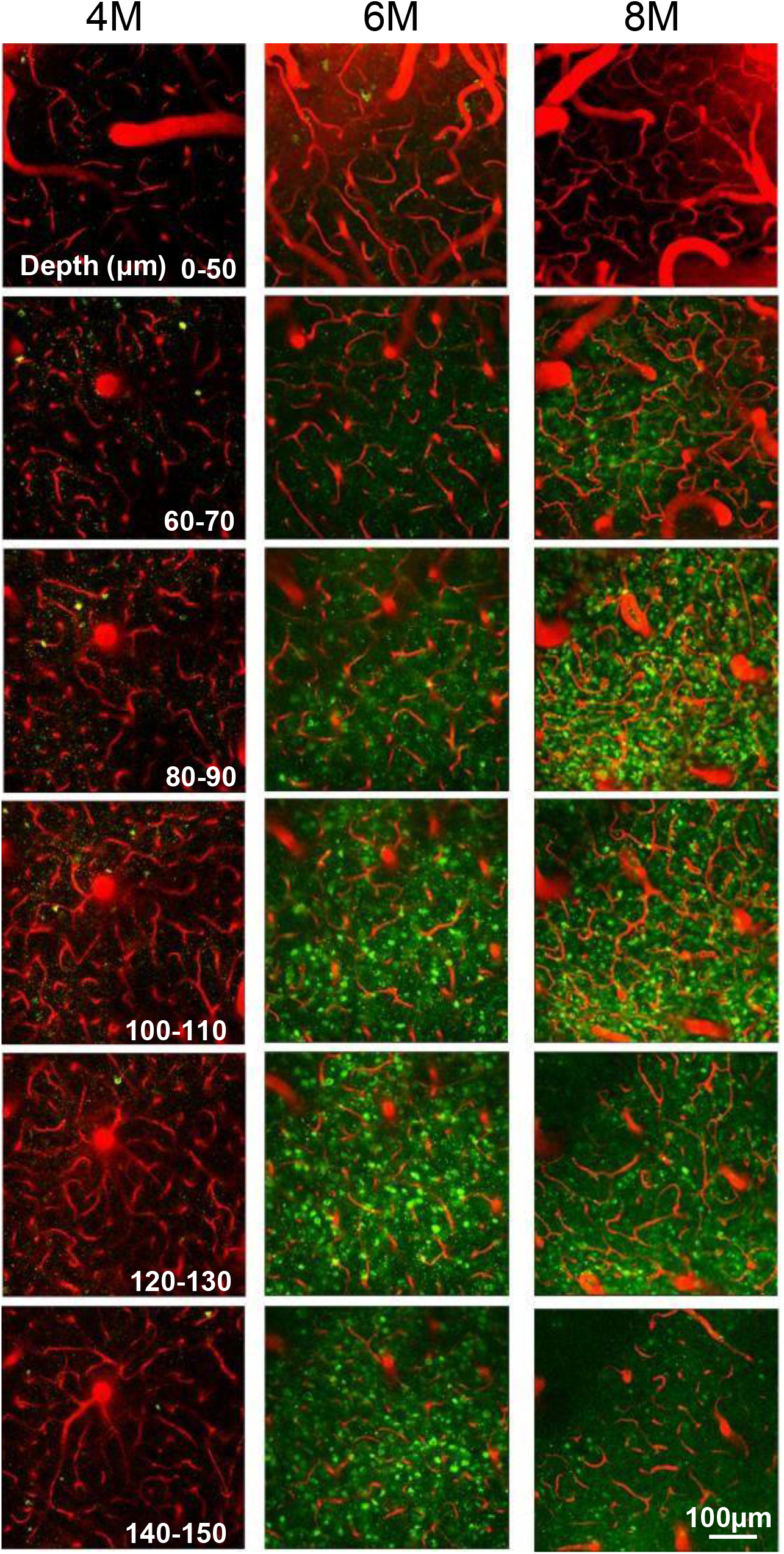
Intravital two-photon laser microscopic imaging of neuronal tau inclusions (green) and blood vessels (red) at different depths of the neocortex of rTg4510 mice at 4 (left column), 6 (middle column) and 8 (right column) months of age. Depths of the focal planes from the brain surface are indicated in left panels. Images were captured at 30 min after intravenous injection of PBB3. Vessels were labeled with SR101.

**Supplemental Figure 2.**
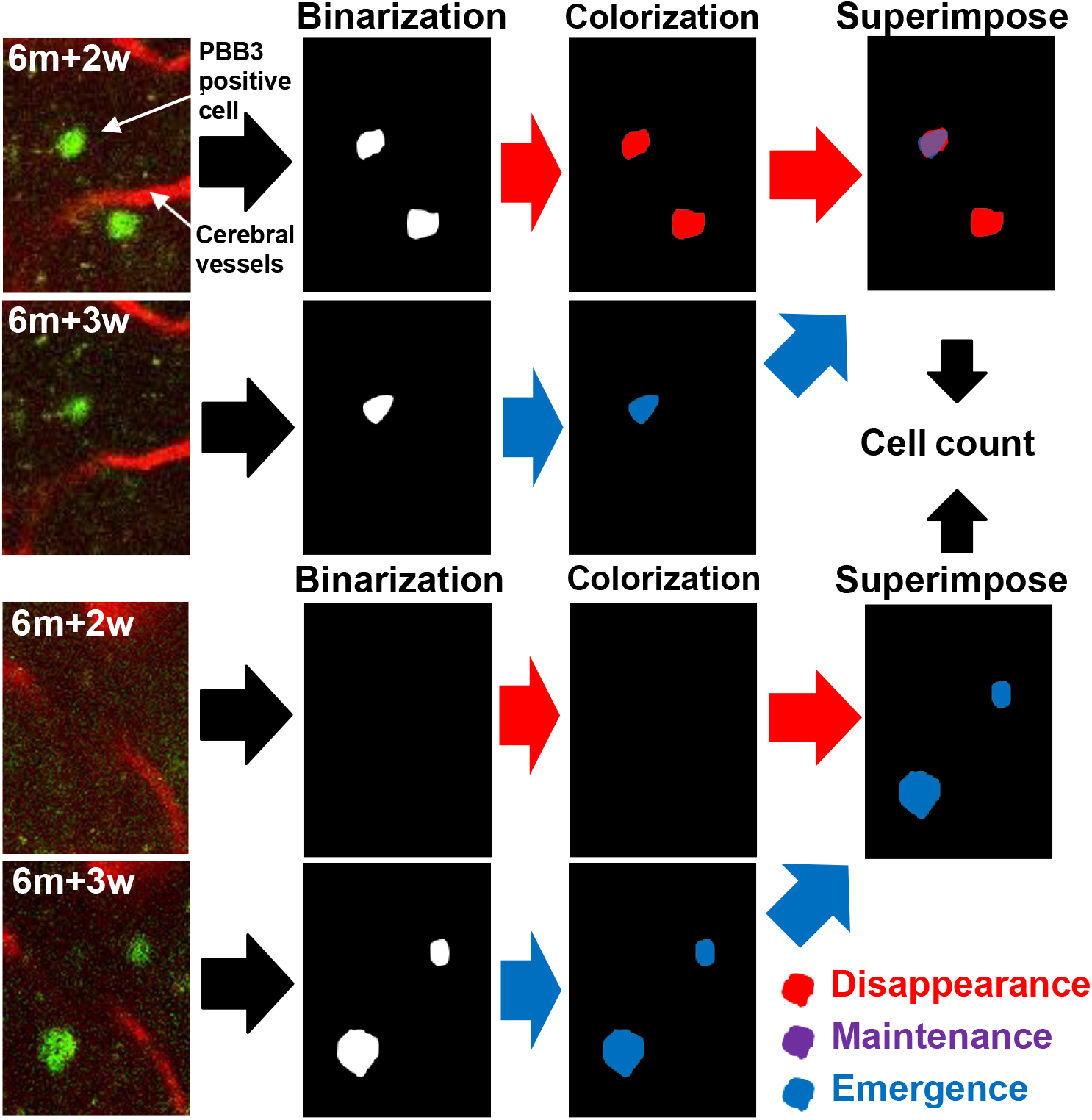
Methods for counting the emergence and disappearance of PBB3-positive tau aggregates during the period of longitudinal two-photon microscopic observations. PBB3 signals were binarized, and signals at baseline and follow-up were pseudo-colored in red and blue, respectively. Merging baseline and follow-up images highlights the emergence and disappearance of aggregates in blue and red, respectively, while persistent presence of aggregates is indicated in purple.

**Supplemental Figure 3.**
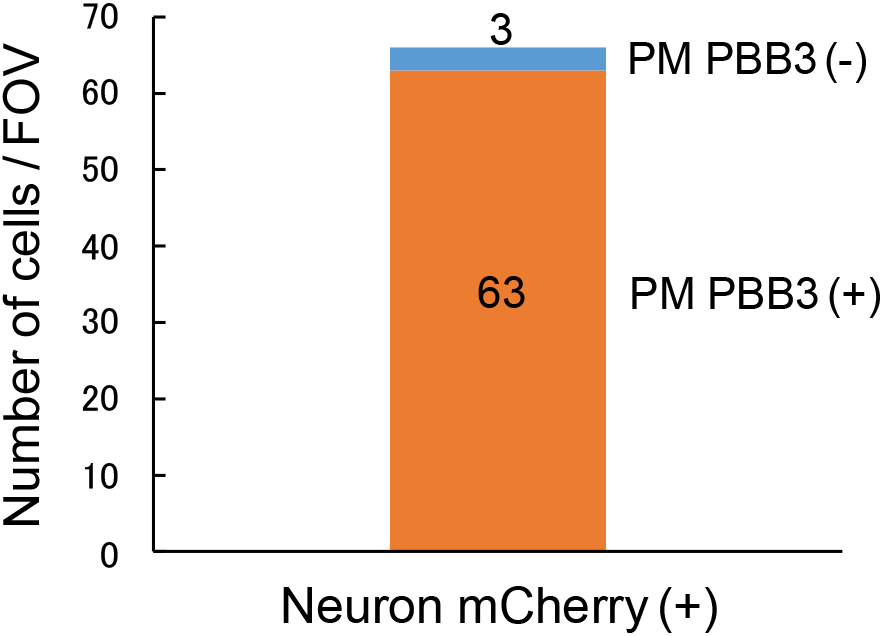
Numbers of mCherry-positive neurons labeled and unlabeled with intravenously administered PM-PBB3 per field of view (FOV) in the neocortex of rTg4510 mice at 6 months of age.

**Supplemental Figure 4.**
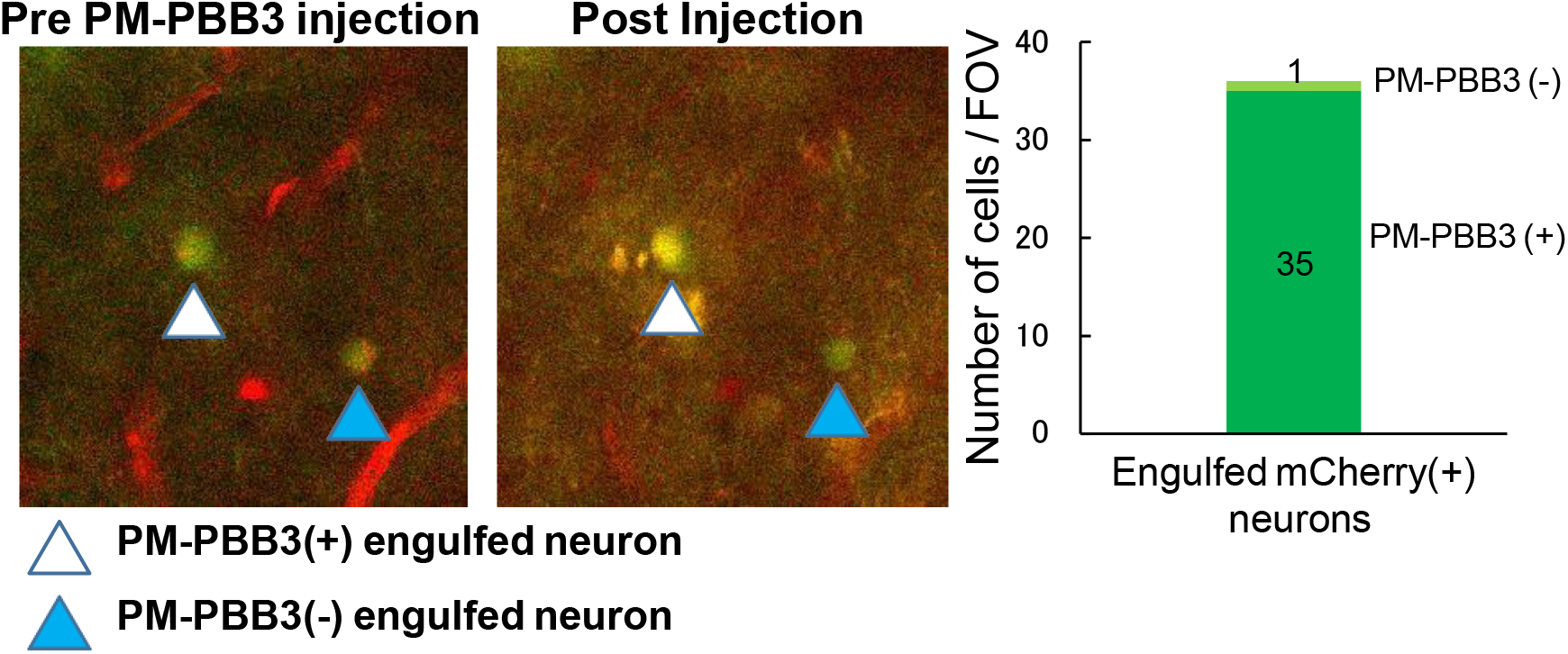
mCherry-positive neurons (arrowheads) engulfed by GFP-positive microglia captured by two-photon microscopy before (left) and 30 min after (right) intravenous injection of PM-PBB3 in the neocortex of rTg4510 mice at 6 months of age. A neuron indicated by white arrowhead was additionally illuminated by PM-PBB3, while fluorescence intensity of a neuron indicated by blue arrowhead was not noticeably altered by the tracer injection. A graph (right) illustrates the proportion of engulfed neurons labeled and unlabeled with injected PM-PBB3 in the neocortical FOV. The actual numbers of PM-PBB3-positive and PM-PBB3-negative neurons are displayed in and above the stacked column, respectively.

**Supplemental Figure 5.**
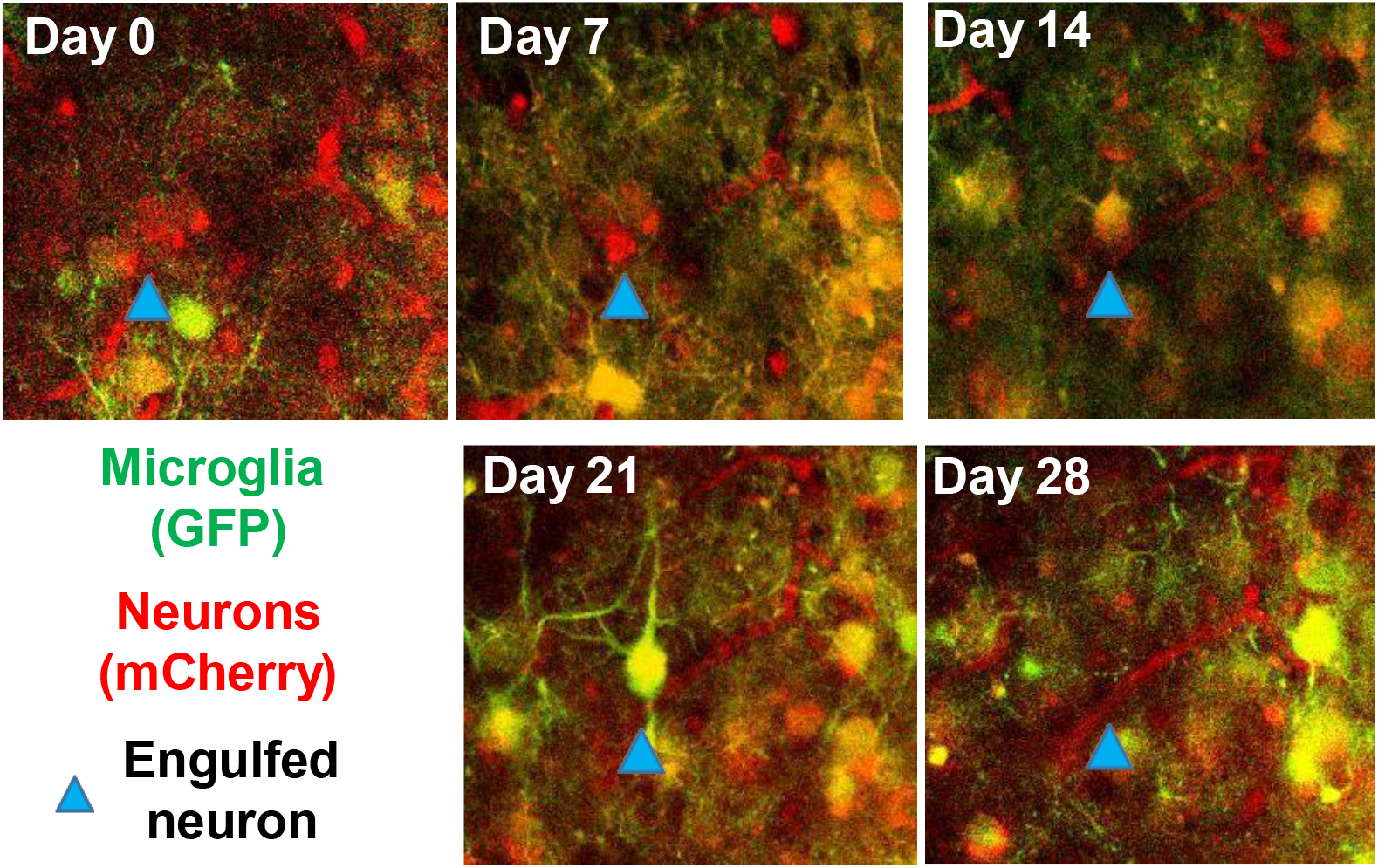
Two-photon microscopic tracking of a neuron engulfed and subsequently eliminated by phagocytotic microglium in the neocortex of an rTg4510 mouse at 6 months of age. A mCherry-positive neuron indicated by arrowhead was not in close contact to microglia at baseline (Day 0) and Day 7, but underwent engulfment by GFP-positive microglia at Days 14 and 21, resulting in phagocytic clearance of this neuron by Day 28. Blood vessels were also labeled in red with SR101.

**Supplemental Figure 6.**
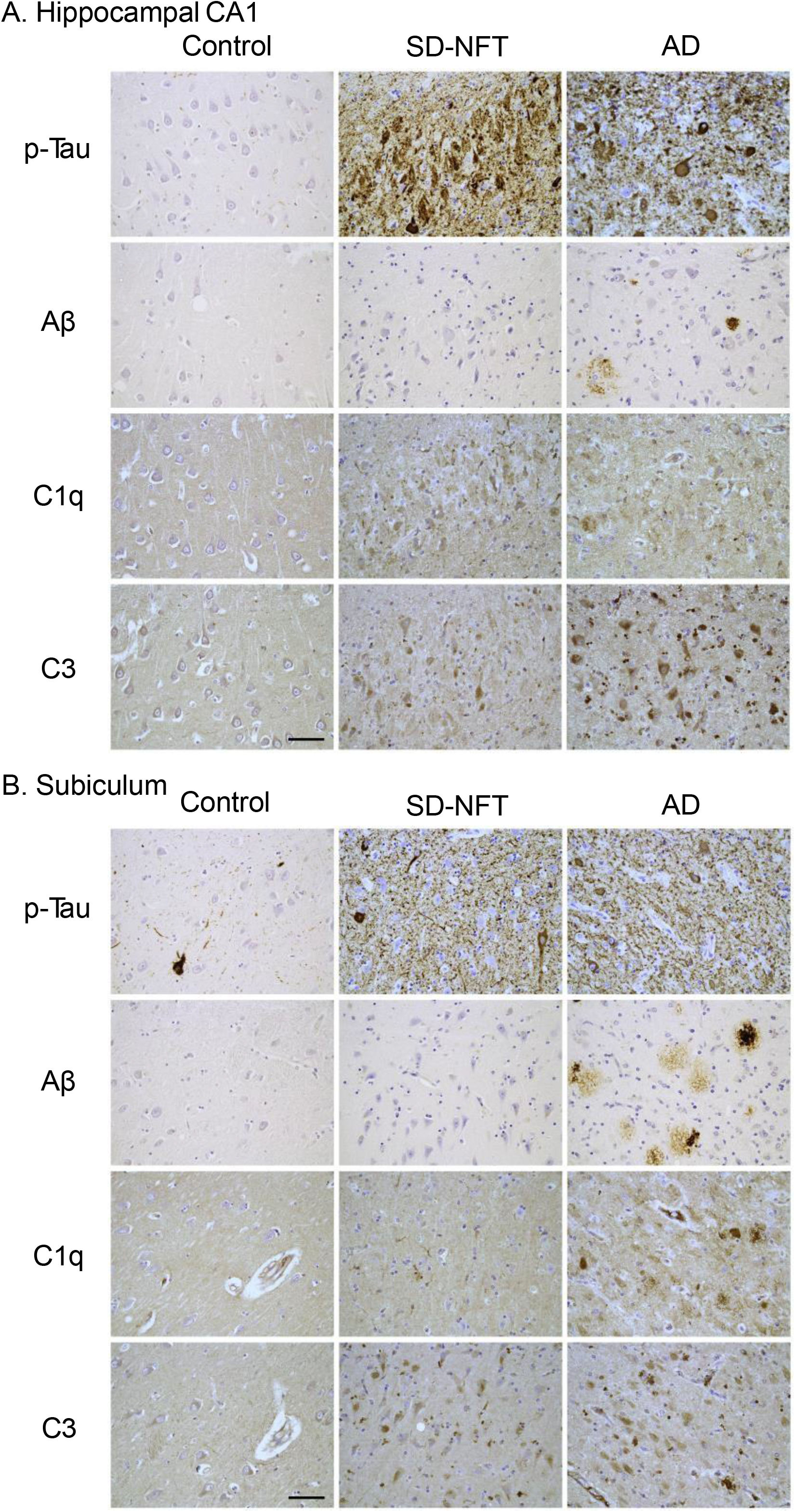
Immunohistochemical staining of phosphorylated tau (p-tau; top row), Aβ (upper middle row), C1q (lower middle row) and C3 (bottom row) in hippocampal CA1 (composite A) and subiculum (composite B) sections derived from an elderly control (left column) and patients with SD-NFT (middle column) and AD (right column). Antibody labeling was developed by 3,3’-diaminobenzidine.

**Supplemental Figure 7.**
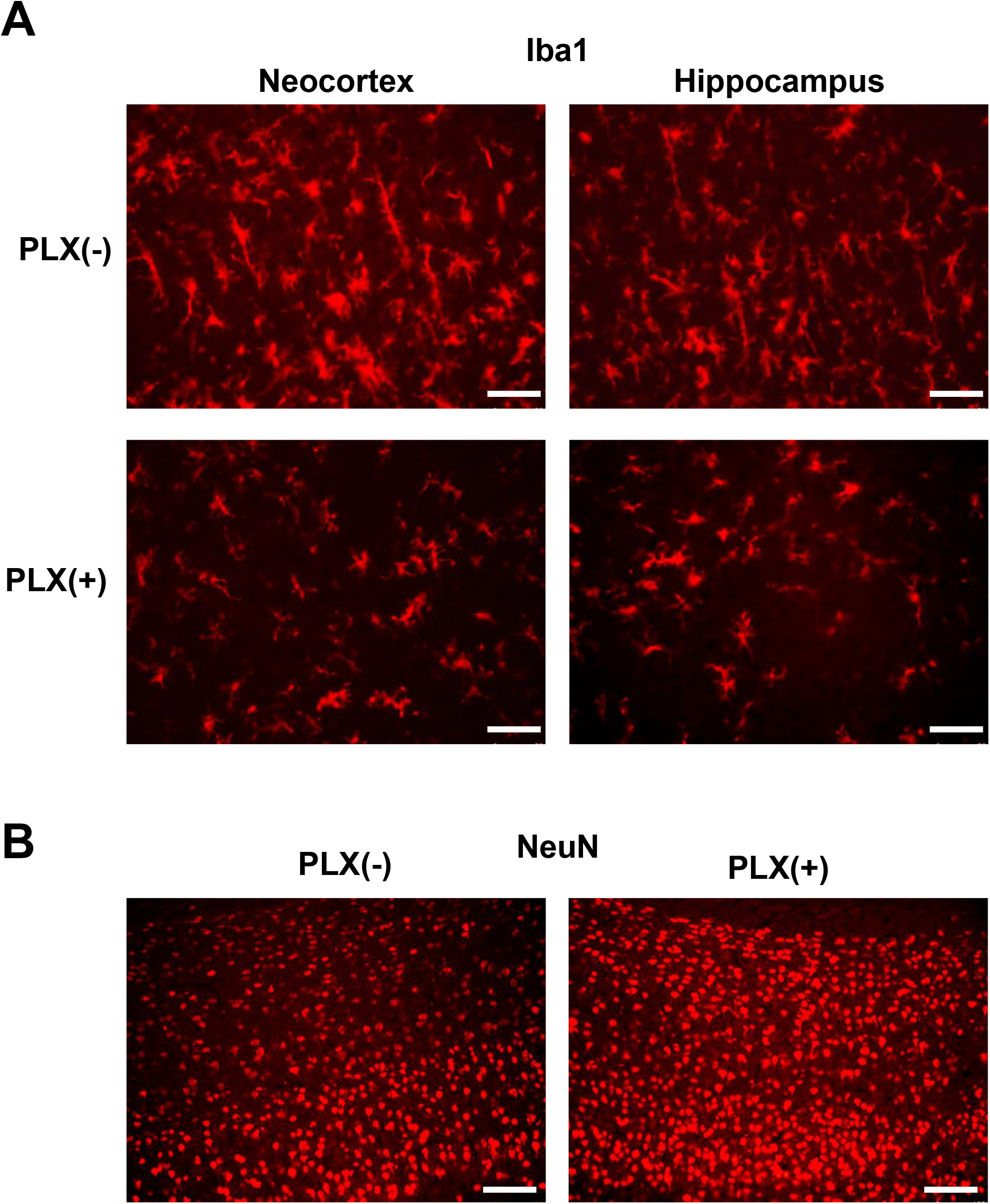
Deactivation and depopulation of microglia and the rescue of neurons in the brains of rTg4510 mice following the PLX treatment. A) Immunofluorescence staining of Iba-1 in brain sections derived from rTg4510 mice. Brain tissues were obtained from 8-month-old mice without treatment [PLX(-)] or following PLX treatment over 1 month [PLX(+)]. Numerous hypertrophic and rod-shaped microglia were noted in the untreated condition, in contrast to a smaller number of ramified and slightly hypertrophic microglia remained after the PLX treatment. B) Immunofluorescence staining of NeuN in neocortical sections sub-adjacent to those displayed in A. Neuronal somas were spared from pathological removals by the PLX treatment. Scale bars, 50 μm (A), and 100 μm (B).

**Supplemental Figure 8.**
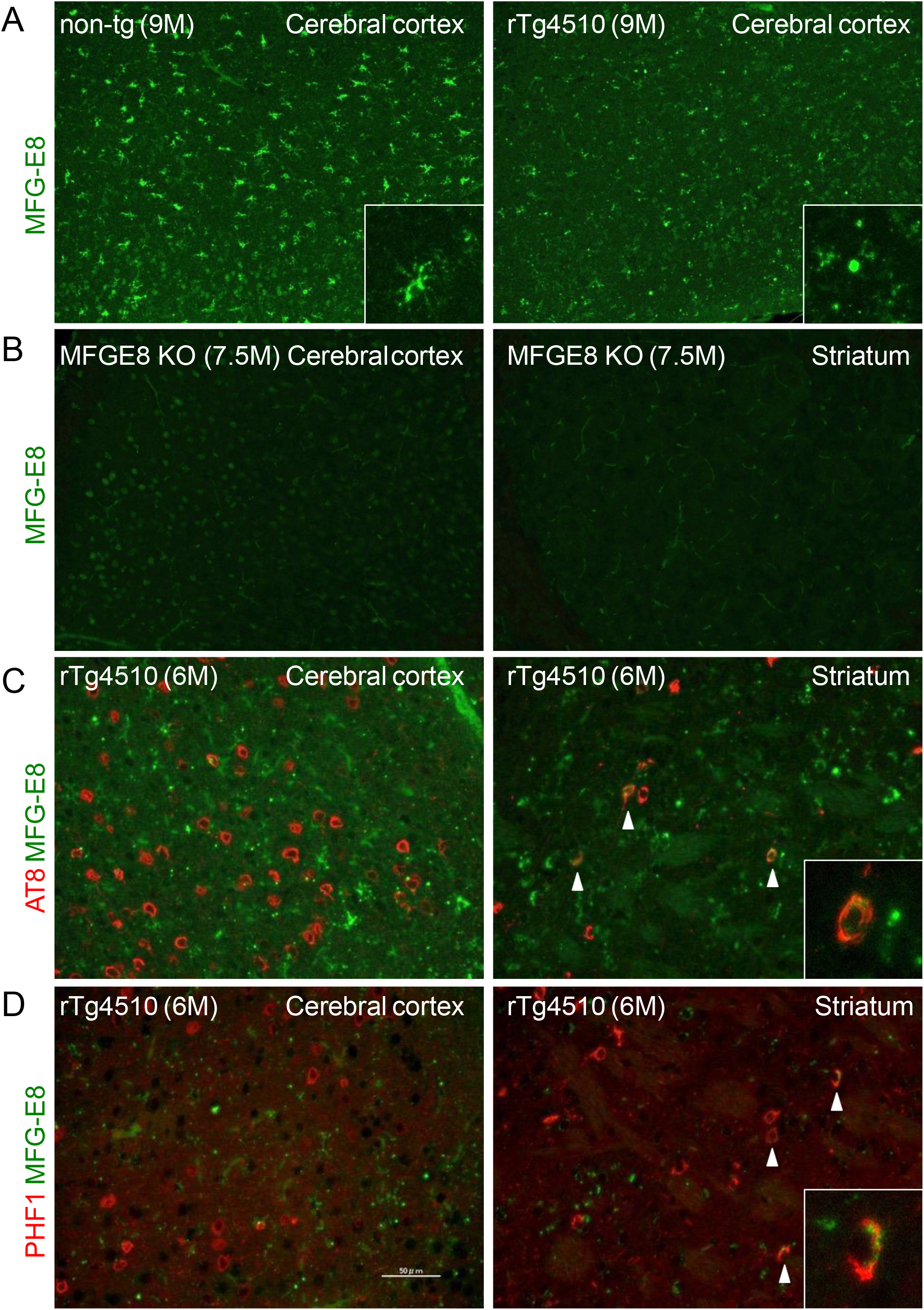
Immunofluorescence staining of MFG-E8 and phosphorylated tau. A) MFG-E8 immunolabeling images in the neocortex of 9-month-old non-Tg (left) and 9-month-old rTg4510 (right) mice. Inbox in the left panel showed glial cell shapes labeled by the MFG-E8 antibody. Inbox in the right panel showed punctate shapes labeled by the MFG-E8 antibody. B) MFG-E8 immunolabeling images in the neocortex (left) and the striatum (right) of a 7.5-month-old MFG-E8 KO mouse. C) Double immunostaining of MFG-E8 (green) and phosphorylated tau (red, AT8 antibody) in the neocortex (left) and the striatum (right) of a 6-month-old rTg4510 mouse. D) Double immunostaining of MFG-E8 (green) and phosphorylated tau (red, PHF1 antibody) in the neocortex (left) and the striatum (right) of a 6-month-old rTg4510 mouse. Co-labeling of MFG-E8/AT8 or MFG-E8/PHF1 was observed in the striatum (arrowheads, inboxes in right panels of C and D). Scale bar, 50 μm.

**Supplemental Figure 9.**
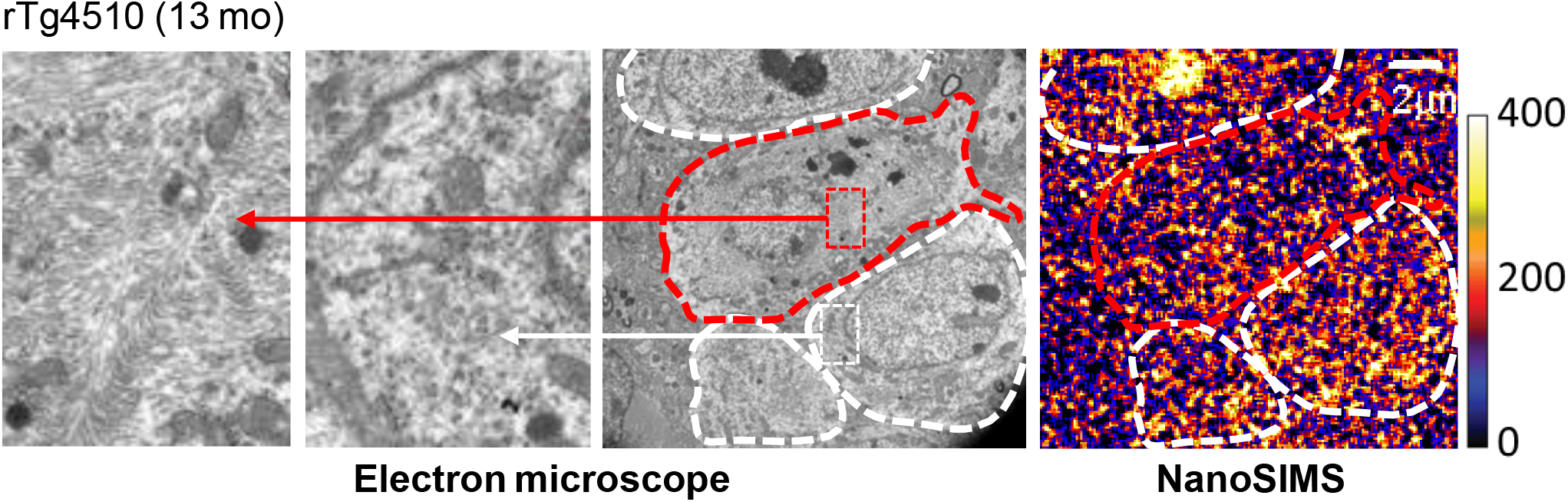
Adjacent ultra-thin neocortical sections derived from an rTg4510 mouse at 13 months of age examined by transmission electron microscope (left three panels) and NanoSIMS (right panel). Neuronal somas containing massive and no visible fibrils putatively corresponding to PBB3-positive tau inclusions are outlined by red and white dashed lines. High-power views of fibril-positive and fibril-negative somas are displayed in left and left middle panels, respectively. NanoSIMS image indicates distributions of signals arising from systemically administered [^13^C]glucose and its metabolites. The color scale denotes arbitrary signal intensity.

## Notes

### Competing Interest Statement

The authors have declared no competing interest.

