## Supplementary material for "Tracking tau fibrillogenesis and consequent primary phagocytosis of neurons mediated by microglia in a living tauopathy model": ONLINE METHODS

Methods, including statements of data availability and any associated accession codes and references, are available in the online version of the paper.

**Animal preparation**

rTg4510 mice (38-48 g, N = 30) and WT mice were used in four separate measurements: 1) animal PET, 2) animal 7T-MRI, 3) two-photon microscopy and 4) NanoSIMS. The parental mutant human tau responder line, the parent tTA activator line, and the resultant F1 rTg4510 mice and littermates were generated and maintained as previously described (Santacruz et al. 2005). Strain backgrounds were maintained with FVB/NJcl (CLEA Japan, Inc. Japan) and 129+Ter/SvJcl (CLEA Japan), respectively. Mice were maintained on a standard diet lacking doxycycline (DOX) to ensure that transgenic human tau was expressed unless the experimental groups with DOX treatment. When the treatment started, mice were given 1.5 g DOX (Sigma Cat# 9891, St. Louis, MO) in 4% sucrose water for 2 weeks. A DOX diet containing 0.2 g/kg DOX (Cat# D13110902, Research Diets INC. New Brunswick, NJ) was administered to mice ad libitum to suppress transgene expression. The animals were maintained and handled in accordance with recommendations of the Guide for the Care and Use of Laboratory Animals and institutional guidelines at the National Institute of Radiological Sciences. All animal experiments were approved by the Animal Ethics Committee of the National Institute of Radiological Sciences. These mice were housed with *ad libitum* food and water in their cages at 25°C in a 12-hr light/dark cycle. All experiments were performed in accordance with the institutional guidelines on humane care and use of laboratory animals and were approved by the Institutional Committee for Animal Experimentation.

**In Vivo PET Imaging**

PET scans were performed using a microPET Focus 220 animal scanner (Siemens Medical Solutions) immediately after intravenous injection of [^11^C]PBB3 (29.7 ± 9.3 MBq). Micro-PET Focus 220 is designed for rodents and small monkeys, providing 95 transaxial planes 0.815 mm (center-to-center) apart, a 19.0-cm transaxial field-of-view (FOV), and a 7.6-cm axial FOV ^1^. Radiosynthesis of [^11^C]PBB3 was described in a previous report ^2^. Before PET measurements, a 31-gauge needle with catheter was inserted into the mouse tail vein for injection of [^11^C]PBB3. After a transmission scan for attenuation correction, a bolus of [^11^C]PBB3 was injected, and an emission scan in 3D list mode was carried out for 90 min. All list-mode data were sorted into 3D sonograms, and were then Fourier-rebinned into 2D sonograms (frames: 1 min x 4, 2 min x 8, 5 min x 14). Images were reconstructed with a filtered back projection using a Hanning filter with a Nyquist cutoff of 0.5 cycle/pixel. Statistical analyses were performed to compare AUC ratio among four mouse groups by two-way analysis of variance followed by Tukey’s test.

**Two-photon imaging in awake animals**

For the surgery procedure, the animals were anesthetized with a mixture of air, oxygen, and isoflurane (3–5% for induction and 2% for surgery) via a facemask, and a cranial window (3-4 mm in diameter) was attached over the left somatosensory cortex, centered at 1.8mm caudal and 2.5mm lateral from the bregma, according to the ‘Seylaz-Tomita method’ ^3^. A custom metal plate was affixed to the skull with a 7-mm-diameter hole centered over the cranial window. The method for preparing the chronic cranial window was previously reported in detail by Takuwa et al ^4,5^. All experiments were performed two weeks after the cranial window surgery.

PBB3 shares features of both tau-binding PET tracer and fluorescent imaging agent^2^. Vessels and abnormal tau protein were fluorescently-leveled with sulforhodamine 101 (SR101; MP Biomedicals, Irvine, CA) and PBB3, respectively. SR101 was dissolved in saline (10 mM) and PBB3 was dissolved in 0.05% DMSO solution, and both fluorescent materials were injected intraperitoneally (SR101: 8 µL/g body weight, PBB3: 2 µg/g body weight) just before initiation of the imaging experiments. The awake animal was placed on a custom-made apparatus, and real-time imaging was conducted by two-photon laser scanning microscopy (TCS-SP5 MP, Leica Microsystems GmbH, Wetzlar, Germany) with an excitation wave-length of 900 nm. An emission signal was separated by a beam splitter (560/10 nm) and simultaneously detected with a band-pass filter for SR101 (610/75 nm) and PBB3 (525/50 nm). A single image plane consisted of 1024 x 1024 pixels, and in-plane pixel size was 0.25–0.45 µm depending on the instrumental zoom factor. Volume images were acquired up to a maximum depth of 0.4–0.8 mm from the cortical surface with a z-step size of 2.5 µm.

Two-photon imaging was performed in awake mice. The experimental protocol for measurements using awake mice was reported previously^4,5^. Briefly, the metal plate on the animal's head was screwed to a custom-made stereotactic apparatus. The animal was then placed on a styrofoam ball that was floating using a stream of air. This allowed the animal to exercise freely on the ball while the animal's head was fixed to the apparatus.

**MRI experiments**

All MRI experiments were performed on a 7.0T horizontal MRI scanner (Magnet: Kobelco and JASTEC, Japan; Console: Bruker Biospin, Germany), with a volume coil for transmission (Bruker Biospin) and a 2-ch phased array surface coil for reception (Rapid Biomedical, Germany). The mice were initially anesthetized with 3.0% isoflurane (Escain, Mylan Japan, Japan), and then anesthetized with 1.5 ~ 2.0% isoflurane and 1:5 oxygen/ room-air-mixture during the MRI experiments. Rectal temperature was continuously monitored using an optical fiber thermometer (FOT-M, FISO, Canada) and maintained at 36.5±0.5˚C using a heating pad (Rapid Biomedical) and warm air. The surface coil was placed over the cranial window (cerebellar cortex). The first imaging slices were carefully set at the rhinal fissure with reference to the mouse brain atlas^6^.

T2-weighted MRI: Transaxial T2-weighted fast spin-echo MR images were acquired using rapid acquisition with a relaxation enhancement (RARE) sequence in the same slice position as the T1-weighted MRI. The imaging parameters were as follows: TR / effective TE = 4,200 / 36 ms, Fat-Sup = on, NA = 4, RARE factor = 8, number of slices = 13, and scan time = 6 min 43 s. Frequency selective saturation pulses and crusher magnetic field gradients were used for fat suppression.

**Adeno-Associated Virus (AAV) Injection**

Full-length cDNAs encoding EGFP and mCherry were amplified by polymerase chain reaction and subcloned into the multi-cloning site of AAV transfer plasmids containing either rat synapsin promoter or human CD 68 promoter with woodchuck posttranscriptional regulatory element (WPRE), and poly-A signal flanked with ITRs. For large-scale preparation of recombinant AAV, AAV plasmid and AAV serotype DJ packaging plasmids (pHelper and pRC-DJ) were introduced into HEK293T cells with polyethyleneimine transfection. 48h after transfection, cells were harvested, lysed, and AAV particles were subsequently purified with HiTrap heparin column (GE healthcare) as described previously ^7^.

**Electron microscope (EM) and NanoSIMS**

A 1.1 M aqueous [U-^13^C] glucose solution (99% ^13^C isotopic enrichment, Sigma-Aldrich, Japan) was prepared for injections into awake mice. A series of doses corresponding to 1 mg/g each were delivered at seven time points by intraperitoneal (i.p.) injections. Mice were euthanized at t = 180 min (injections at 0, 10, 20, 60, 90, 120, and 150 min).

Transmission EM was performed on mice after they were deeply anesthetized and sacrificed by transcardial perfusion with 5 ml of saline followed by 30 ml of fixative (2.5% glutaraldehyde, 2% paraformaldehyde, 0.05% calcium chloride in 0.1 M cacodylate buffer). Samples were prepared and examined with JEOL 1200EX EM.

After EM imaging, thin sections were mounted on 10-mm diameter, aluminum mounts with double-sided copper sticking tape, and coated with a 10-nm layer of gold. The distribution of C isotopes (i.e. ^13^C/^12^C ratio) was determined in the same areas with a NanoSIMS 50L instrument following established procedures ^8,9^. NanoSIMS images were processed with L’IMAGE© software; count-rate smoothing (3 by 3 pixels) was systematically applied to all images. By overlaying a semi-transparent NanoSIMS image over an electron micrograph of the same region, it is possible to assign individual pixels in the NanoSIMS image to specific cell types or their sub-cellular compartments. Quantified δ^13^C enrichments can therefore be ascribed to each structure. Neurons in layer IV were imaged by NanoSIMS. From the EM images, neurons with/without tau were identified according to their morphology.

**Histological examination**

The animals were deeply anesthetized with sodium pentobarbital and transcardially perfused with saline, and brain tissues were removed. The brains of rTg4510, PS19, MFG-E8 knockout mice and non-Tg littermates were immediately frozen with dry ice for C3 immunostaining. For C1q immunostaining, the brains of these animals were fixed with 4% paraformaldehyde in phosphate buffer and cryoprotected using 30% sucrose in phosphate buffer. Twenty-µm-thick frozen sections were generated in a cryostat (HM560; Thermo Fisher Scientific). Fresh frozen sections were post-fixed in 4% paraformaldehyde solution. The brain sections were immunostainined with rabbit monoclonal antibody against C1q (1 : 1000, Abcam, ab182451), mouse monoclonal antibody against synapsin (1 : 2000, Synaptic Systems, 106011), tau phosphorylated at Ser202 and Thr205 (AT8, 1 : 250, Thermo Fisher Scientific, MN1020), rabbit polyclonal antibody against C3 (1 : 1000, DAKO, A0062), mouse monoclonal antibody against Iba-1 (1 : 1000, Merck Millipore, MABN92), mouse monoclonal antibody against NeuN (1 : 1000, Merck Millipore, MAB377), and goat polyclonal antibody against MFG-E8 (1:500, R&D systems, AF2805). Fluorescence images were captured using DM4000 (Leica) and BZ-X710 (Keyence) microscopes.

The use of human autopsy brains was approved by the Human Ethics Committee of Fukushimura Hospital, and written informed consent for the postmortem analysis was obtained from each subject or their relatives. Formalin-fixed brains underwent systematic and standardized sampling, and neuropathological evaluation by a single experienced neuropathologist (H.Y.). Amyloid histopathology was assessed according to CERAD criteria ^10^. NFT pathology was staged according to Braak and Braak ^11^. Diagnoses were reached on the basis of clinical history and NIA-Regan criteria. Cases were diagnosed for a high likelihood of AD using NIA-Regan criteria. Cases of senile dementia of the neurofibrillary tangle type (SD-NFT), a subset of dementia characterized by numerous NFTs in the hippocampal region and absence of amyloid plaques throughout the brain, were diagnosed according to criteria as described elsewhere ^12-14^. Paraffin-embedded sections (4 μm thick) of temporal cortex including hippocampal region from 3 patients with AD, 3 patients with SD-NFT, and 3 non-neurological controls were used in this study. Immunohistochemistry was performed on the sections using the Histofine Simple Stain Max PO (Multi) kit (Nichirei, Tokyo, Japan), which is an amino acid polymer labeled by peroxidase and goat anti-mouse and anti-rabbit IgG Fab’. An anti-human Aβ (mouse monoclonal 6F/3D, 1:50 dilution, Vector laboratories, Burlingame, CA), an anti-phospho-tau (mouse monoclonal AT8, 1:1,000 dilution, Innogenetics, Gent, Belgium), anti-C1q (mouse monoclonal 9A7, 1:2,000 dilution, Abcam, Cambrige, UK) and anti-C3 (mouse monoclonal B-9, 1:50 dilution, Santa Cruz, Dallas, TX) were used for primary antibodies. To activate antigen, the rehydrated sections were incubated with formic acid for staining of NFTs and amyloid plaques, or incubated with 10 mM citric acid in a pressure cooker for C1q and C3 immuno-staining. Endogenous peroxidases were inactivated by treatment with 0.3% H_2_O_2_ in distilled water. The sections were incubated with primary antibody at 4°C overnight and then with anti-mouse and anti-rabbit immune-peroxidase polymer (Nichirei) for 30 min at 25°C. Peroxidase activity was detected with chromogen/substrate reagent containing 3,3’-diaminobenzidine (Nichirei). Sections were counterstained with hematoxylin and dehydrated. Images were captured with a Leica microscope (DM4000, Lecia, Wetzlar, Germany) equipped with a color digital camera.

**Treatment of mice with a TSPO ligand, Ro5-4864**

Six month-old rTg4510 mice (n=5) were injected (i.p.) with TSPO- ligand Ro5-4864 (3 mg/kg; 2.5 % DMSO, 1% tween in saline) (Sigma Aldrich, C5174). In order to evaluate the mobility and phagocytosis of microglia, two-photon microscopy imaging of microglia, neurons and vessels was started 2 weeks before Ro5-4864 injection and measurements were made twice before and twice after injection. Microglial mobility and phagocytosis were measured by comparing the positions of microglia and neurons in the two images taken either before or after injection. Microglial movement is recorded when the microglia position changes with respect to the cerebral blood vessels in one week (Fig. 6A, white triangle). Similarly, microglia phagocytosis occurs when a neuron surrounded by microglial foot processes disappears within one week (Fig. 6B (bottom), white triangles).

**Treatment of mice with a CSF1R inhibitor, PLX-3397**

PLX-3397 (PLX) was formulated in standard chow by Research Diets Inc. at the dose of 290 mg/kg. Unlike a previous report on the treatment of inbred C57BL/6 and hybrid C57BL/6/129 mice ^15^, the PLX administration to rTg4510 mice on a hybrid FVB/129 background induced partial depletion of microglia in the brain. We started feeding 4-month-old rTg4510 mice chow with or without PLX and volumetric MRI scans of these animals were conducted at 6 months of age. Similarly, feeding of food with or without PLX to 7-month-old rTg4510 mice was initiated and the treatment was terminated at 8 months of age. The brains of these animals were subsequently collected for immunohistochemical examinations.
